# Preprocessing of general stenotic vascular flow data for topological data analysis

**DOI:** 10.1101/2021.01.07.425693

**Authors:** Christopher L. Bresten, Jihoon Kweon, Xinjuan Chen, Young-Hak Kim, Jae-Hun Jung

## Abstract

A new analysis and classification method of vascular disease based on topological data analysis (TDA) has been proposed in [1]. The proposed method utilizes the application of persistent homology to hemodynamic variables. Particularly, 2D homology is obtained from the velocity field of the flow projected onto the unit sphere, known as so-called the *S*^2^ projection. It was shown that such homology is closely related to the degree of vascular disease. The original method was developed based on the computational fluid dynamic (CFD) solutions of the straight stenotic vessels.

In this paper, we develop a preprocessing method that enables the proposed TDA method to be applied to general stenotic vessels of irregular geometry. The velocity field is subject to a coordinate transformation correcting for orientation and curved geometry. The preprocessed data is projected onto *S*^2^ and the corresponding homology is calculated. We show that this preprocessing is necessary for the proposed TDA method to be successfully applied to general types of stenotic vessels. Validation was performed on a set of clinical data including reconstructed vascular geometry with corresponding diagnostic indices.

## Introduction

Cardiovascular disease is the leading cause of death worldwide. Accurate diagnosis and prediction is crucial. The diagnosis methods today are based on the anatomical approach by investigating the geometry of the diseased vessel using image analysis and the functional approach by measuring the pressure drops across the stenosis. The gold standard diagnostic method uses fractional flow reserve (FFR) values, which are calculated by dividing a distal pressure measurement by a pressure reading proximal to the patient’s stenosis [2, 3]. Newer approaches utilize computational fluid dynamics (CFD). Patient-specific CFD is also a possible analysis tool [4] to estimate FFR based on the simulated data. Recently, machine learning approaches were also proposed as an alternative diagnosis methodology [5, 6, 7]. For these methods, clinicians need a canonical index that can represent the collective knowledge of the diseased vessel. Especially for the machine learning approach, one needs to extract features of the vascular disease for the given patient’s data.

In [1] a new method of providing a measure of vascular disease has been proposed based on topological data analysis (TDA) [8]. In order to apply TDA to vascular flows for the proposed method in [1], the velocity field was projected onto a unit sphere (*S*^2^), referred as the *S*^2^ projection. It was shown that this simple projection method is highly efficient and powerful for visualizing and quantifying the vascular disease, with which the flow complexity is measured by observing the directional information of the velocity fields. For example, the overall flow pattern is laminar for the healthy vessel and is more turbulent when there is stenosis. On *S*^2^, the laminar flow is projected as a point cloud centered around one pole on the sphere while the fully developed turbulent flow provides a point cloud all over the sphere including two poles. Then the proposed TDA method computes the persistent homology of the point cloud on *S*^2^. It was shown that the two dimensional homology of *S*^2^ projection of vascular flows is closely related to the degree of disease and can be used along with the value of FFR for diagnosis. The original development of the proposed method was based on the 3D spectral computational fluid dynamic (CFD) data of the regular straight stenotic vasculature.

In this paper, we verify that the proposed TDA method can be applied effectively to curved vessels with more general irregularity in a similar way to the case of the straight vessel. In order to apply TDA to general vessel type, we develop a projection method which preprocesses the hemodynamic data such that it becomes suitable for TDA. The proposed projection algorithm is based on the singular value decomposition (SVD) and principle component analysis (PCA). After the preprocessing, we decompose each vessel into small segments and TDA is performed to the velocity fields on *S*^2^ in all the segments. We use the real patient MRI data provided by Asan Medical Center, Seoul, Korea. The numerical results clearly show that the persistent homology after the preprocessing projection is significantly correlated to the FFR values, hence can be used a meaningful tool for the prediction of cardiovascular disease.

## Methods and materials

### Clinical data

Two invasive angiographic images were selected at the end of the diastolic phase of the cardiac cycle, then combined to build three dimensional geometry using CAAS Workstation 7.5 (Pie medical Imaging, Maastricht, NL). This was done in compliance with the Declaration of Helsinki and approval from the IRB of Asan Medical Center was granted by waiver of research consent.

### CFD simulation

The boundaries of the vasculature were assumed rigid and discretized using Ansys ICEM CFD 15.0 (Ansys Inc, Canonsburg, PA, USA). The vascualr flow was modeled as an incompressible fluid with density 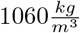 and Newtonian 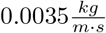. The inflow rate was set 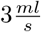 at and outflow boundaries on the opposite face of the vessel set. The steady state solution was solved for using Ansys Fluent 15.0. Bifurcated vascular geometry was not included. A first order finite element method was used with a mixed mesh of tetrahedral and hexahedral elements.

### TDA Software

For the TDA computations JavaPlex [9] was used through its MATLAB interface. The Lazy witness complex was used, with landmark points selected by max-min selector; 75 landmark points were used. Pre-processing was done in Python.

### Notation

In this paper, we let matrices be denoted in capital letters, and indexed using subscripts, so that *X_i,j_* represents the *ij*th element of the matrix *X*. Also we let the subscript in this context mean an entire row or column, so that *X_∗,j_* will denote the *j^th^* column and vice-versa. For the given *X*, let *X^H^* denote the Hermitian (conjugate transpose) of *X*.

For the hemodynamic variables, let *ρ* = *ρ*(**x***, t*) be the density, *p* = *p*(**x***, t*) the pressure, **u**= (*u, v, w*)^*T*^ the velocity vector for the position vector **x**= (*x*^0^*, x*^1^*, x*^2^)^*T*^ ∈ Ω and time *t* ∈ ℝ^+^. Here Ω is the closed domain in ℝ^3^.

### Preprocessing

The preprocessing happens in several steps. First the orientation of the vessel is regularized by PCA on the spatial coordinate point cloud. Then the vessel is segmented along the first principal axis, followed by an iterative process of segmentation and fitting of a cubic spline. This results in a curvilinear coordinate system that reflects the curvature of the vessel, which can be used to project the velocity field and segment the vessel.

The process assumes that the geometry of the vessel is contiguous, non-bifurcating, and has high aspect ratio.

Let ***x**_k_* be the coordinate vector, 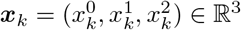 where the solutions to the incompressible Navier-Stokes equations are calculated and *X* ∈ ℝ^*n*×3^ be the set composed of all 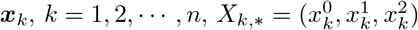. ***x**_k_* constitute the point cloud we use for the analysis. Here 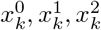 denote the *x*-, *y*- and *z*-coordinate of each point cloud. To each ***x_k_***, the hemodynamic variables are assigned, e.g. the 3D velocity vector 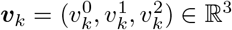 and the pressure *p_k_* ∈ ℝ^+^. ***v**_k_* and *p_k_* are all functions of ***x**_k_* and *t*. We define *W* ∈ ℝ^*n*×3^ as the velocity field, composed of ***v***_*k*_ such that 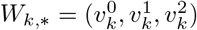.

In order to discover this principle axis, effectively regularizing orientation, we employ PCA. The principle axis is the direction of highest aspect ratio, along which the blood flows. Despite the local and global curvatures of the vessel, the principle direction should unidirectional and this direction will be mapped into a straight line along the principle axis. To perform PCA, we first compute the mean coordinate values for each dimension of ***x***_k_, *μ*_0_*, μ*_1_ and *μ*_2_ such that

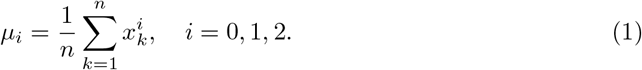

Let *T*_1_ be the affine transformation that translates *X* by *μ* as below

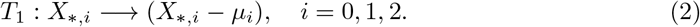

The principle axis can then be found using the SVD. In the following SVD plays a crucial role. Each point in space can be thought of as a sample of a three dimensional random variable in our setting. After centering the data, the matrix *T*_1_(*X*) *T*_1_(*X*)^*T*^ is then the covariance matrix of the sampled random variable, which projects a linear combination to a vector of covariances. The eigenvectors of this operator are the linear combinations which give a multiple of themselves as covariances. These eigenvectors can be interpreted as the directions in the space where the sample of the random variable has the most variation. These eigenvectors are the right singular vectors of *T*_1_(*X*). This gives us a basis that we can use to represent the data, the one corresponding to the largest singular value having the largest covariance, and therefore the accounting for the more variation in the data than any other possible choice of basis vector. Since the points are a point cloud defining the geometry of a three dimensional object, this gives us the direction where the object is longest, effectively orienting the geometry in the way that is ideal linear approximation for us to chop it up in the direction of the flow field.

Let the following be the SVD of *T*_1_(*X*)

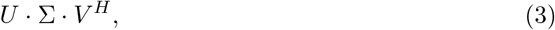

where *U* ∈ ℝ^*n*×*n*^ and *V* ∈ ℝ^3×3^ are unitary matrices and Σ ∈ R^*n*×3^ is a rectangular diagonal matrix with non-negative real values on its diagonal. Since *T*_1_(*X*) is real, *U* and *V* are orthonormal, i.e. *U^H^* = *U* and *V ^H^* = *V*. The principle axes/directions are the columns of *V*. Let *T*_2_ be the linear transformation of *X* projected onto the right principle vectors *V*

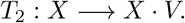

Let 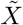 be the following

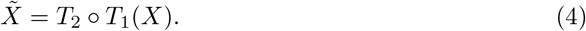

In practice, one can use a sparse sampling of *X* for the SVD as it also provides a good approximation to *V*. This is the case particularly because we are dealing with only 3 dimensions while there is a large number of points, so the shape of the vessel is well represented by a relatively sparse random samples. Figure 1 shows the schematic images of *T*_2_ ◦ *T*_1_. The original vessel in finite element mesh domain (left figure) is mapped into the new domain along the principle axes through *T*_2_ (right figure). The colored intervals are the segments of the vessel (see below for the domain decomposition).

**Fig 1.**
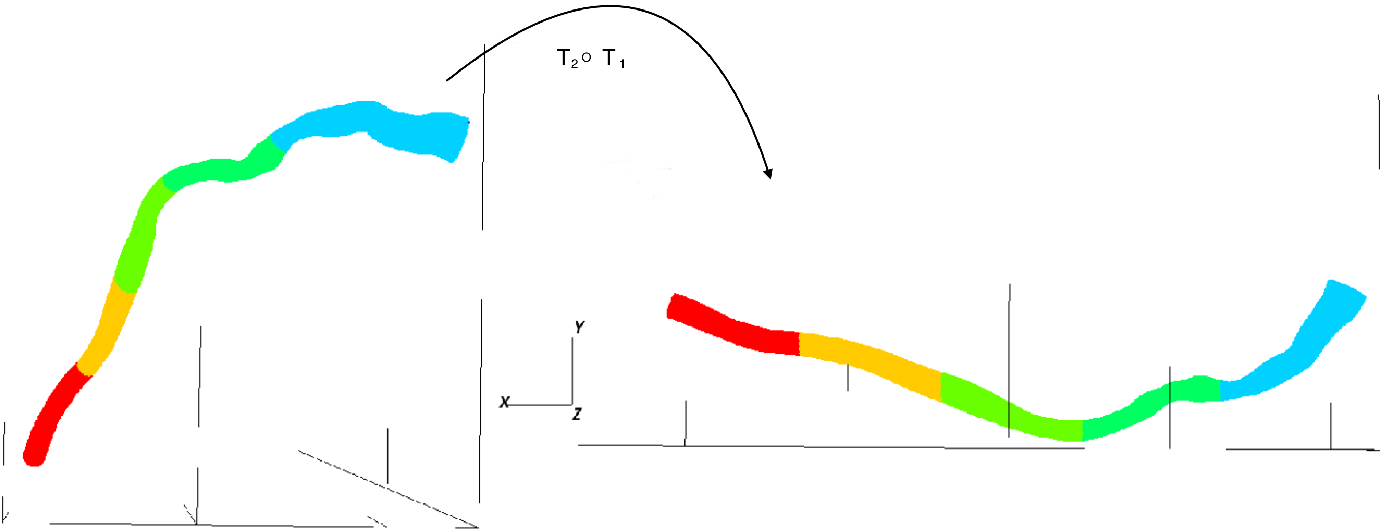
Orienting the vessel. Coordinate transformation *T*_2_ ◦ *T*_1_ of *X*. The original coordinates of the vessel (left) are transformed into the new coordinate along the major axis (right). The colors show the initial five segments made along the principal axis.

Now we want to partition the vessel into *m* parts along the principal axis. First we find the maximum coordinate and minimum coordinate values in the first principle direction,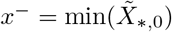 and 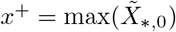. We split the vessel into *m* non-overlapping intervals by diving the total interval 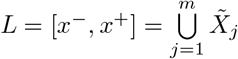. Let *X_j_* be the *j^th^* segment corresponding to 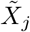 in the original coordinate system. Similarly *W_j_* is the corresponding velocity data corresponding to *X_j_*.

## Center line projection

The process above results in a set of initial conditions to a nonlinear parametric fit to the vascular geometry, which will be used to project the velocity field. The next step, is to begin the nonlinear fit by calculating the centroids of the segments *X_j_*:

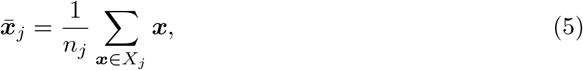

where *n_j_* is the number of elements in *X_j_*. Now with *m* centroids, we can fit a parametric curve through them. The fitting can be done exactly in a collocation sense, but we choose cubic splines with a small smoothing parameter to keep the derivative continuous. We will denote this as ****P**** (*t*), where *t* ∈ [0, 1] is the parametric variable. We first use this curve ****P**** to re-partition the vessel into a larger number of points *M*, this is done by nearest neighbors to each of the *M* points on the curve.

The centroids are calculated again, and a new curve is fit to these centroids. This can be done several times to get a better fit. The following Figure 2 shows the resulting bins after several iterations, with a final value of *M* = 80:

**Fig 2.**
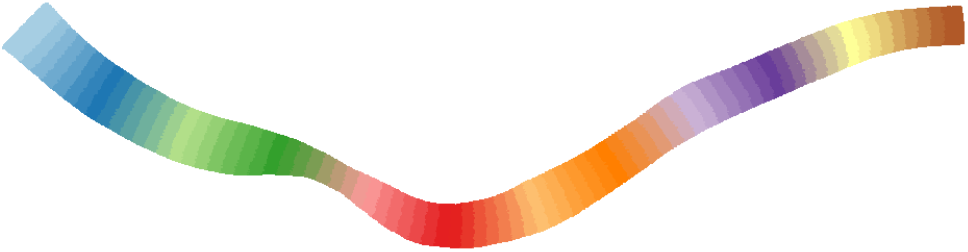
Segmented vessel. Partitioning of the vessel into 80 segments along cubic spline fit. Each color is a different segment.

Now that we have a good curve fit, we proceed to construct the projection operator for each point in *X*. This is done by sampling both ****P**** and 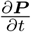 at a larger number of points *N_p_* = 300, *t_i_ ∈ T* ⊂ [0, 1]. This sampling is the one used to project the velocity field. Then, like the above segmentation process, we solve the nearest neighbor problem for each ****x**** ∈ *X*:

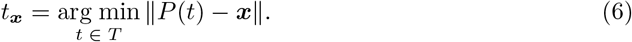

*t_**x**_* allows us to address the curve ****P**** (*t*) for each point ****x****. The normalized derivative is then calculated:

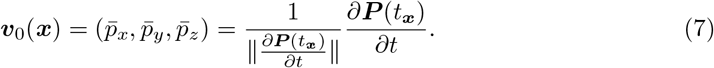

Now, we find the null space of this vector and call them ****ν****_1_(****x****) and ****ν****_2_(****x****). Then we stack them into a projection matrix

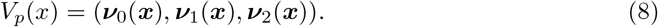

This is used to project the velocity vector coresponding to point ****x****. This is done for all ***v*** ∈ *W*.

## Algorithm

In this section, we summarize the preprocessing algorithm and provide the pseudo codes. The algorithm is broken into several functions and several stages for clarity. The first stage finds the longest axis along the geometry and partitions the data along this axis. This is a linear approximation to seed the process. The centroids of each partition are used as points to fit a parametric cubic spline. This cubic spline is then used to partition the geometry into more segments, again calculating the centroids, and fitting another spline.

After several rounds of this, the cubic spline fit is used as the centerline, approximating the first dimensional geometry of the vessel. This is used in a coordinate transform on the velocity field. The result is a velocity field represented in terms of its flow with respect to the geometry of the vessel. Figure 3 shows the velocity fields before (left) and after (right) the curvilinear projection. Notice how the sign of the velocity flips due to the curvature of the vessel in the left image (red area), while this is absent in the right image.

**Fig 3.**
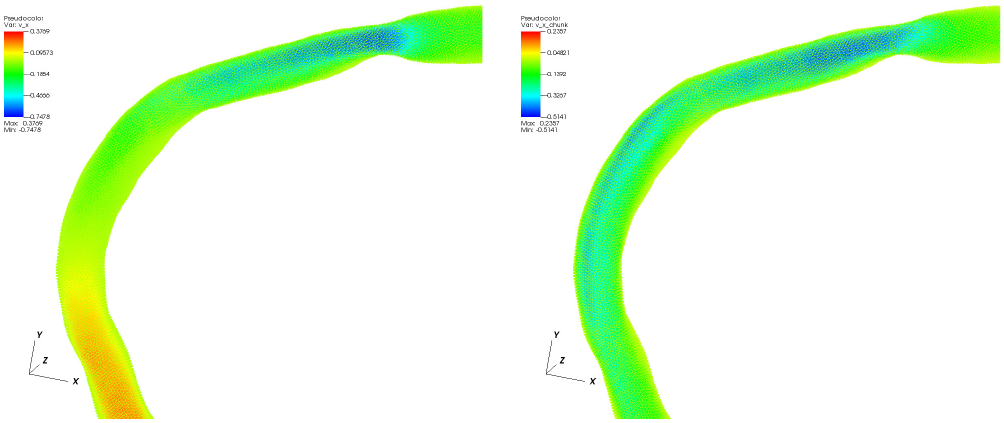
Transformed velocity field. Left: *v*_0_(*x*− direction) before projection, Right: *v*_0_ after curvilinear projection.

**Fig 4.**
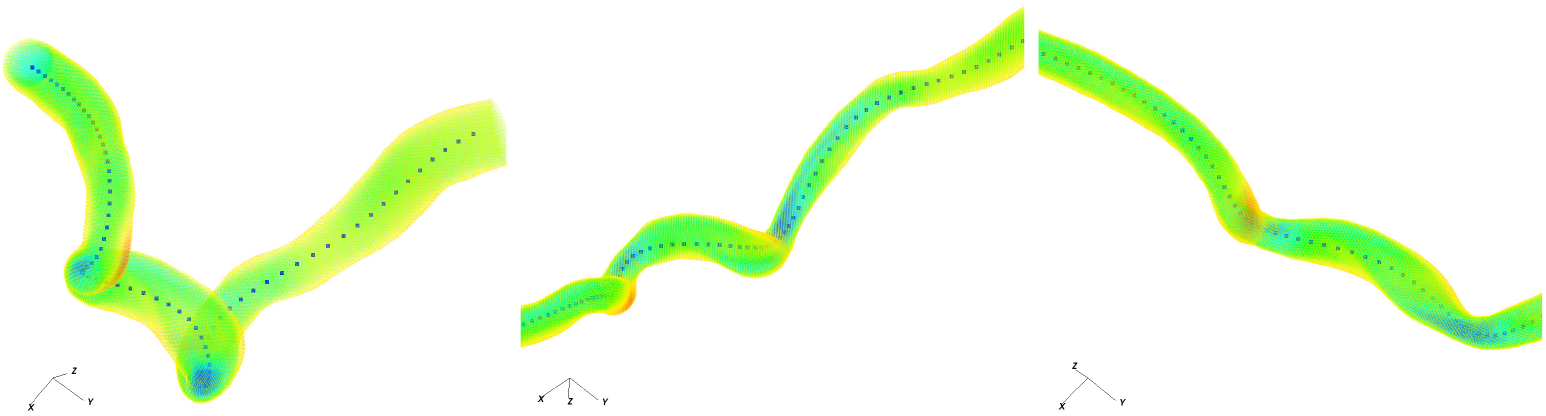
Parametric cubic spline fit. The least-squares fit parametric spline curve plotted to the vessel, also referred to as the centerline.

**Algorithm 1:**
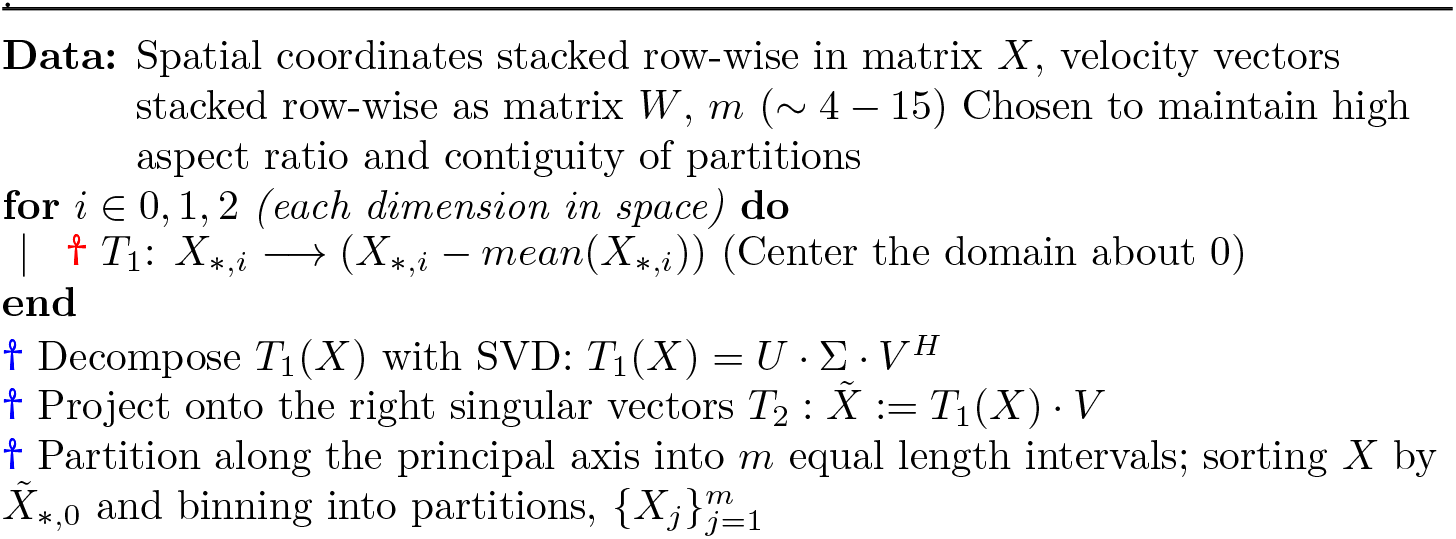
Initial linear partitioning. This serves serves as the initial conditions for the cubic spline fit.

**Algorithm 2:**
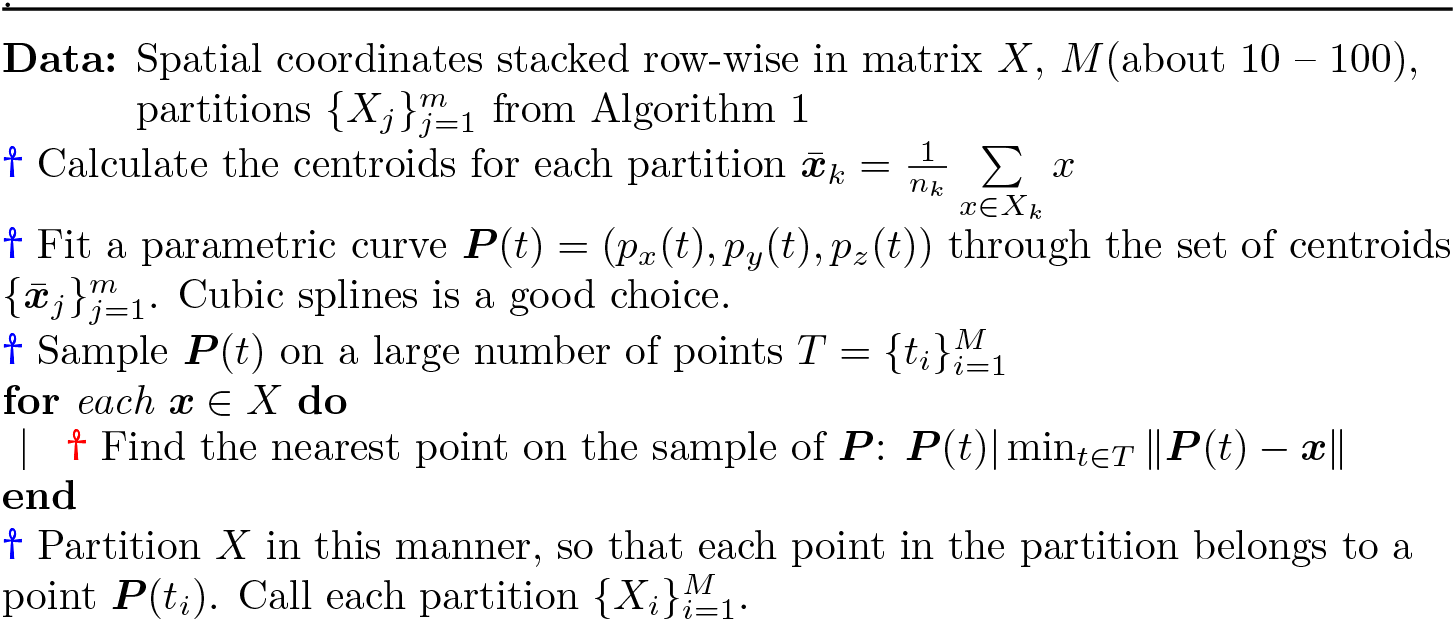
Segmentation. Partitioning along nonlinear axis.

**Algorithm 3:**
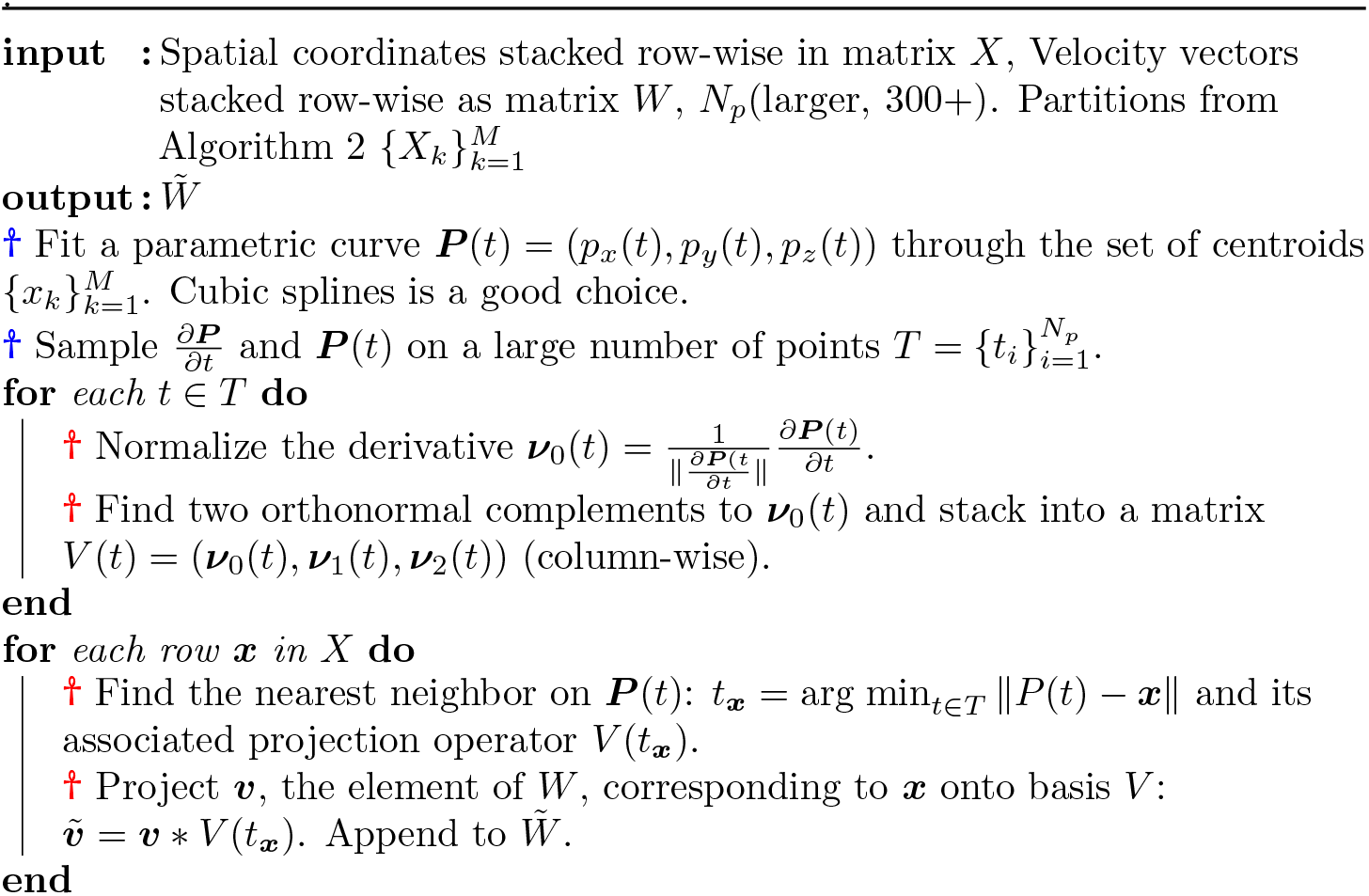
Algorithm 3: Projection of velocity field. Transform of velocity field onto curvilinear co-ordinates.

**Algorithm 4:**
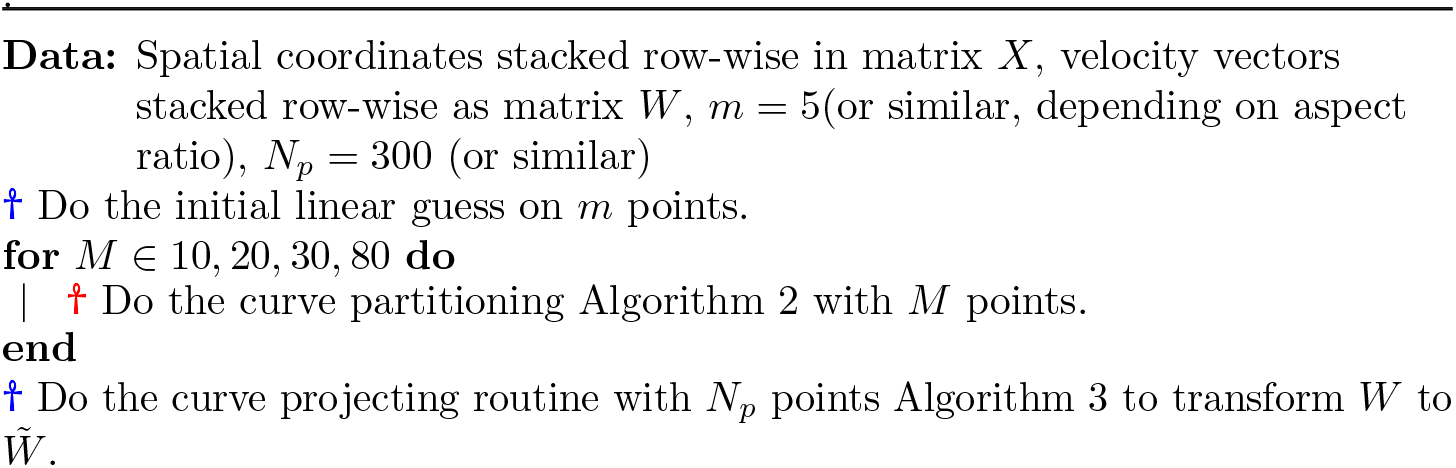
Algorithm 4: Main loop. Main loop connecting all previous components.

## Topological data analysis

Once all the necessary preprocessing and transformations are carried out, we apply TDA to the transformed velocity field. As mentioned in Introduction, the main purpose of this paper is to show the applicability of the TDA analysis method proposed in [1] to clinical data of general type of vasculature and to show that the proposed method is efficient in describing stenotic disease. First we must briefly explain the key element of TDA that we apply to the flow field. For more details, we refer readers to [1, 10].

Let 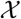 be the topological space. The topological space 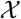 that we consider in this paper is the space constructed by the velocity field. Note that, however, it is possible to construct 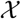 from more hemodynamic variables than the velocity field such as the pressure variable. Although more general space is interesting, such cases are not considered in this paper. In order to construct a meaningful topological structure given the point cloud, we first explain singular homology. Homology is a topological invariant that describes the holes of various dimensions of 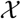.

The assumption is that there exists 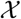 and the point cloud is a set of sampling out of 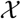. Singular homology is on 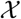 rather than the point cloud. An *n*-simplex is a convex set composed of *n* + 1 vertices (e.g. points in velocity field in our case). For example, 0-simplex is a point, 1-simplex is an edge, 2-simplex is a filled triangle and 3-simplex is a filled tetrahedron. The *n*-simplex is the *n*-dimensional version of the triangle. The standard *n*-simplex is the convex hull of the standard basis in R*n*+1. Singular *n*-simplex in 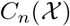 is a continuous map *σ* from the standard *n*-simplex to 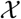. Let 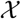 be the free abelian group whose basis is the set of singular *n*-simplices in 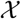. Then we can define the boundary map, *δ_n_* from 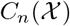 to 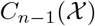 where 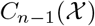 is constructed from 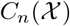 by removing the vertices 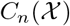 of one at a time. Then the *n*-dimensional homology group *H_n_* is defined by 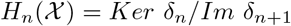 where *Ker* and *Im* are the kernel and image groups, respectively. Roughly we interpret the rank of *H_n_* as the number of holes residing in the *n*-dimension of 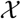. Thus if we can find *H_n_* for *n* = 0, 1, 2,…, we can obtain a rough idea of the structure 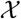 of in terms of holes in it. However, it is difficult to find *H_n_* of 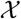 because 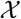 is arbitrary in general. Thus, instead of dealing with the homology of 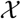, we use rather the homology of the roughly imitated space of 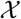.

To imitate the original space 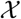 algorithmically, we consider a simplicial complex *K* which is a collection of simplices. The collection satisfies the conditions that *K* contains all lower-dimensional simplices of *σ* if *σ* is a simplex of *K* and that the intersection of two simplices in *K* is a simplex in *K* if the intersection exists. Then in the similar way, the homology group of *K*, *H_n_*(*K*) is defined as 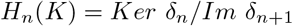 where *δ_n_* is defined similarly as above. *K* is a collection of simplicies, so given the point cloud, we basically connect simplices hierarchically. The number of generators of *H_n_*(*K*) is known as the Betti number, *β_n_*. Roughly speaking, *β*_0_ is the number of the connected components of the resulting *K*, *β*_1_ is the number of the one-dimensional holes of *K*, e.g. the loop or cycle structures of *K*, *β*_2_ is the number of the two-dimensional holes of *K*, and so on.

In this work, we use the Vietoris-Rips simplicial complex, for which we consider the metric space 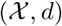 where *d* is a usual Euclidean metric in this paper. Then the Vietoris-Rips complex, *V* (*τ*) is built on the point cloud out of 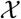 having a *k*-simplex for every collection of *k* + 1 vertices within in a distance *τ* of each other in the Euclidean metric. Here, *τ* is known as the filtration parameter. Thus the Vietoris-Rips complex is constructed with the scale of *τ*. Once constructed, we can compute *H_n_* of *V* (*τ*). However, it is difficult to know which value of *τ* can generate the most similar simplex to 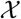. Thus, instead of finding *H_n_* for a fixed value of *τ*, we construct *V* (*τ*) for various values of *τ* in ascending order, say, *V* (*τ_i_*), *i* = 0, 1, 2, · · · and *τ_i_ < τ_j_* for *i < j*. Then we have various *H_n_*(*V* (*τ_i_*)) for various *τ_i_*. Clearly given *N* number of vertices, if *τ* = 0, there are *N* vertices isolated and not-connected and obviously *β*_0_ = *N*, *β*_1_ = 0*, β*_2_ = 0, and so on.

### Persistence Barcode

The variation of homology with respect to *τ* yields the concept of persistent homology. The collection of such homology versus *τ* is known as the barcode. The barcode provides the information of the Betti number and the information of the persistence, the length of the generating barcode, across *τ*. The persistence tells how long the given geometric hole structure maintains with *τ*. The persistence, the interval of each barcode, plays an important role in our method. Particularly, the persistence of the 1D and 2D barcodes are closely related to the degree of flow complexity when the velocity field is mapped onto the unit sphere. The zero-dimensional barcode of *H*_0_ is the graph of *β*_0_ versus *τ*. For the given *N* vertices, *β*_0_ = *N* for *τ* = 0. Thus all the 0-dimensional barcodes of *H*_0_ starts from *τ* = 0 while those barcodes of higher dimension than 0 starts from *τ* where the corresponding hole forms.

The following diagram shows the barcodes for *H*_0_ (left) and *H*_1_ (right), respectively, for the point cloud which is composed of the four vertices of the unit square. From the left barcode we know that there are four vertices given (4 bars in *y*-axis) when *τ* = 0. We also know that when *τ* = 1 all the four vertices are connected and there remains only one connected component afterwards, which is visualized as the single barcode labeled 4 in *y*-axis. As all the four points are connected while no vertices are connected diagonally because *τ* = 1 is smaller than the diagonal distance of two vertices. Thus we know that one hole forms at *τ* = 1 and it lasts until 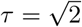 where two vertices are connected diagonally and two triangles form. Note that once triangle is formed, inside is filled as a 2-simplex. Then the hole also disappears, which is the end point of the one-dimensional barcode in the right figure.

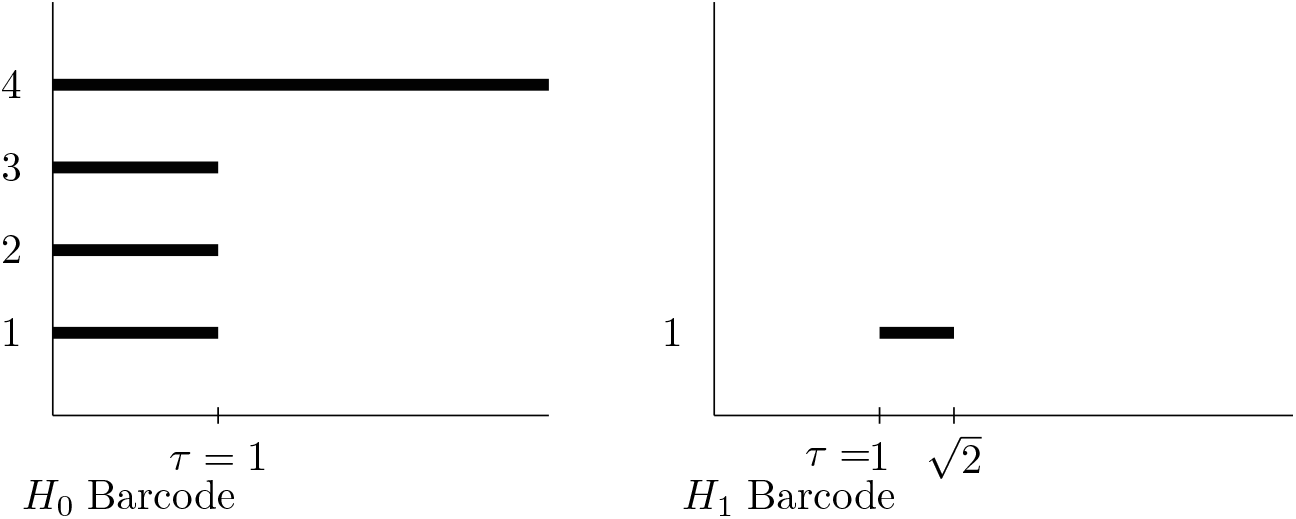

### Persistent homology of vascular flows

TDA of vascular disease is to construct a simplicial complex based on the velocity fields. For the given point cloud (velocity fields in our case), a simplicial complex is created by gluing a finite number of simplices together, for which a gluing condition is needed. For the condition, the topology is described with a metric with the parameter *τ* explained above. The distance between two simplices is determined using the Euclidean metric *d*. For example, two 0-simplices must be connected forming an edge if their Euclidean distance is less than or equal to the value of *τ*. Here we note that it is hard to use the raw data of vascular flows for TDA. In [1] it was shown that the raw data needs to be transformed into a point cloud that is suitable for a meaningful TDA. We use the *S*^2^ projection proposed in [1] also described in the following, but we also note that this is not a unique transform. There could be a better transform for the construction of the simplicial complex.

Figures 5 and 6 show the sample *S*^2^ projection of the curved vessel obtained from the patients’ data. The color represents the pressure value, which is absent from our analysis but helpful in this image. As shown in the figures, the *S*^2^ projection shows how the velocity fields are distributed. It is interesting to observe the existence of the circular pattern on *S*^2^. We hypothesize that those circular patterns are related to the prediction of the disease development, which we do not aim to investigate but will be considered in our future work. In TDA such circular patterns are summarized in 1D homology *H*^1^, while existence of voids due to multidirectional flow are described by *H*_2_.

**Fig 5.**
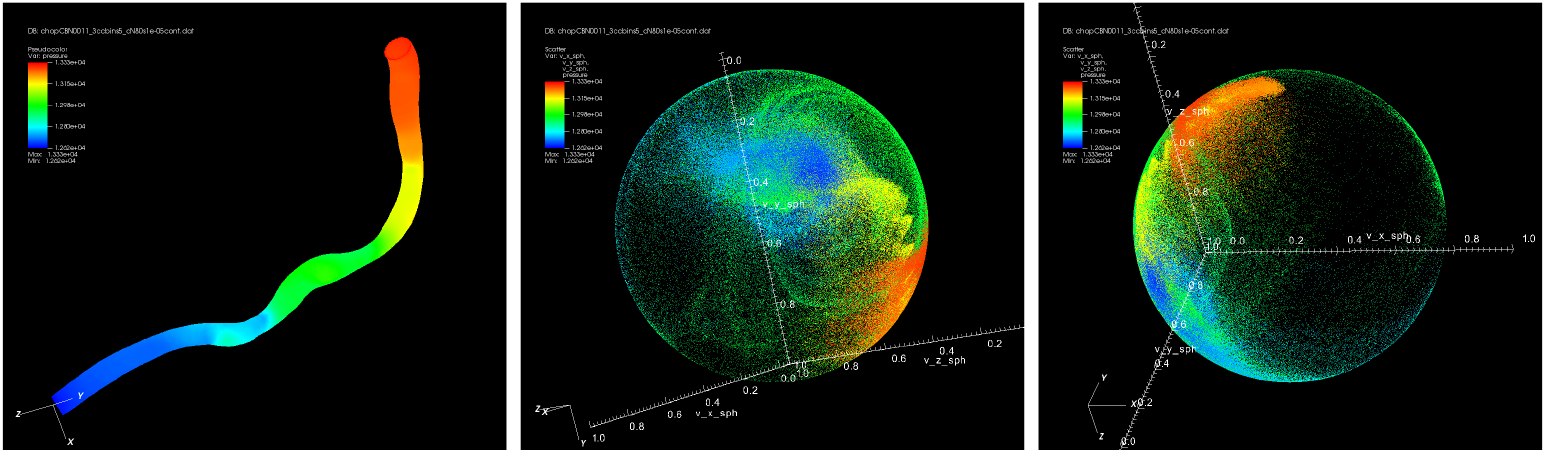
Sphereical projection. Sample spherical projection with pressure as color variable(middle, right), and pseudocolor plot of pressure (left) on the actual morphology of the stenotic vessel.

**Fig 6.**
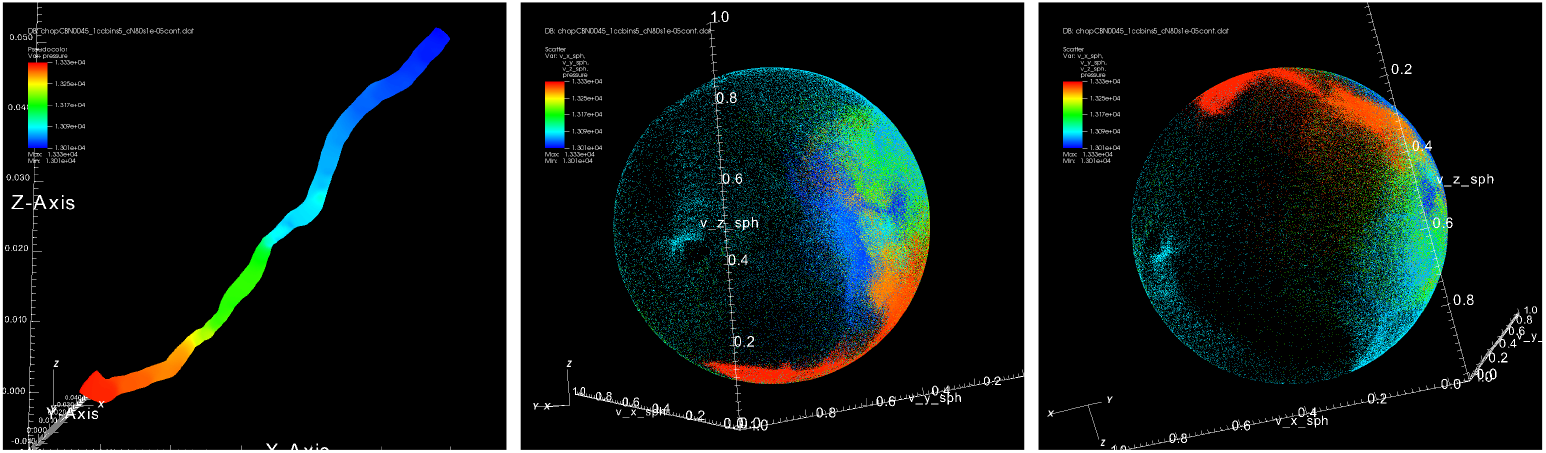
Sphereical projection. Sample spherical projection with pressure as color variable(middle, right), and pseudocolor plot of pressure (left) on the actual morphology of the stenotic vessel.

## Application to the preprocessed vessel

Calculating the persistent homology on the entire vessel is prohibitively expensive, so we perform out TDA on small segments of each vessel, as depicted in Figure 2. Even with the centerline projection, there is a large amount of noise in the boundary regions of the vessel. Due to the spherical projection, this noise appears as any other velocity vector to the TDA steps. Operating on small segments of the vessel helps alleviate inaccuracies caused by this noise.

According to [1], the 2D persistence homology of the velocity fields projected on *S*^2^ can be used to determine whether there is a stenosis. Therefore, by decomposing the whole vessel into smaller segments and calculating the 2D persistences homology within each one, we can get the identity of the segment where stenosis occurs. With the centerline projection methodology described previously, the segmentation of vessels is obtained by chopping up the 3D space of vessels into *M* small segments along with the parametric curve fit. One can see that the size of each segment is not necessarily the same since it depends on the slope of the centerline in the segment region.

We work with the maximum persistence in both *H*_1_ and *H*_2_. This is the length of the longest interval in the persistence diagram, which can be visually understood as the longest line in the bar code. The maximum *H*_2_ persistence is used to determine the segment closest to the stenosis.

## Results

Here we validate the proposed method using clinical data, which is the primary result of this paper. We show a statistically significant correlation between the FFR and the topological persistence calculated in the manner described in the previous sections. To be specific, after segmentation of vessels with preprocessing, we consider the maximum *H*_1_ and *H*_2_ persistences in each segment of vessels, which are the maximal length of the generated 1D and 2D barcodes of the given point cloud, i.e. the velocity field projected on *S*^2^ in this work. We find that there is highly weak correlation in the case where the preprocessing steps are omitted. This is due to the confounding factor introduced by the arbitrary orientation of the vessel in the spatial domain, which results in a misrepresentation of the velocity field, where the direction of the flow changes with respect to the coordinate system, but not with respect to the vessel. Preprocessing with centerline projection alleviates this unwanted factor, and provides an appropriate segmentation of the vessel along its natural curvature.

The results are given in Table 1. The column labeled *Segment number* is the number of the segment in the vessel where the largest maximum *H*_2_ persistence occurs, which is presumably the location of the stenosis. The column labeled *FFR* is the FFR value, which is estimated by dividing a distal pressure by proximal pressure, both of which are measured after a probable lesion is located using X-ray angiography. And the column labeled *H*_1_ *Persistence* is the maximum *H*_1_ persistence in the segment where the largest maximum *H*_2_ persistence occurs. And the column labeled *H*_2_ *Persistence* is the value of the largest maximum *H*_2_ persistence, i.e., the maximum *H*_2_ persistence in the corresponding *Segment*. The first column is the vessel id.

**Table 1.** Results: For each vessel, the FFR value, maximum *H*_1_ and maximum *H*_2_persistences in the segment where the largest maximum *H*_2_ persistence occurs.

| Vessel Id. | FFR | $H_1$ Persistence | $H_2$ Persistence | Segment number |
| --- | --- | --- | --- | --- |
| CBN0759 | 0.62 | 1.01179 | 0.22 | 54 |
| CBN2122 | 0.66 | 0.41905 | 0.3 | 21 |
| CBN0012 | 0.69 | 0.82216 | 0.16 | 17 |
| CBN3355 | 0.69 | 0.53659 | 0.18 | 18 |
| CBN0006 | 0.7 | 1.05928 | 0.14 | 21 |
| CBN1290 | 0.7 | 0.52892 | 0.12 | 21 |
| CBN3222 | 0.7 | 0.71805 | 0.18 | 52 |
| CBN0047 | 0.72 | 0.49718 | 0.18 | 34 |
| CBN3366 | 0.72 | 0.67079 | 0.28 | 22 |
| CBN0045 | 0.73 | 0.46359 | 0.42 | 45 |
| CBN0018 | 0.74 | 0.75791 | 0.3 | 38 |
| CBN1255 | 0.74 | 0.55535 | 0.22 | 36 |
| CBN1353 | 0.75 | 0.64994 | 0.22 | 8 |
| CBN1711 | 0.76 | 0.79459 | 0.16 | 32 |
| CBN3265 | 0.77 | 0.4601 | 0.14 | 12 |
| CBN1960 | 0.78 | 0.77871 | 0.32 | 7 |
| CBN1492 | 0.78 | 0.65907 | 0.12 | 61 |
| CBN3284 | 0.78 | 0.32153 | 0.56 | 33 |
| CBN2034 | 0.79 | 0.90161 | 0.18 | 49 |
| CBN0754 | 0.79 | 0.64229 | 0.14 | 33 |
| CBN1368 | 0.8 | 0.87848 | 0.1 | 37 |
| CBN1203 | 0.8 | 0.33451 | 0.16 | 71 |
| CBN0874 | 0.8 | 0.39672 | 0.52 | 27 |
| CBN1741 | 0.8 | 0.79987 | 0.2 | 50 |
| CBN0569 | 0.8 | 0.60789 | 0.36 | 39 |
| CBN1088 | 0.81 | 0.60526 | 0.14 | 18 |
| CBN2043 | 0.81 | 0.72963 | 0.24 | 53 |
| CBN3254 | 0.81 | 1.1454 | 0.12 | 36 |
| CBN0022 | 0.81 | 0.2421 | 0.5 | 30 |
| CBN1185 | 0.81 | 0.45952 | 0.2 | 70 |
| CBN1345 | 0.82 | 0.49201 | 0.38 | 32 |
| CBN1510 | 0.82 | 0.34708 | 0.44 | 3 |
| CBN3400 | 0.82 | 0.35935 | 0.14 | 42 |
| CBN1103 | 0.83 | 0.60585 | 0.12 | 39 |
| CBN1484 | 0.83 | 1.01112 | 0.1 | 50 |
| CBN0252 | 0.83 | 0.38384 | 0.38 | 51 |
| CBN1084 | 0.84 | 1.24371 | 0.12 | 41 |
| CBN1089 | 0.84 | 0.38813 | 0.26 | 48 |
| CBN1868 | 0.85 | 0.35258 | 0.38 | 44 |
| CBN0003 | 0.85 | 0.46665 | 0.12 | 49 |
| CBN0540 | 0.85 | 0.719 | 0.24 | 41 |
| CBN0014 | 0.85 | 0.30592 | 0.44 | 41 |
| CBN1419 | 0.86 | 0.34468 | 0.26 | 28 |
| CBN0918 | 0.86 | 0.41882 | 0.2 | 65 |
| CBN1731 | 0.86 | 0.59057 | 0.16 | 79 |
| CBN0460 | 0.86 | 0.43247 | 0.44 | 45 |
| CBN1245 | 0.87 | 0.34222 | 0.52 | 39 |
| CBN0011 | 0.87 | 0.37225 | 0.2 | 41 |
| CBN0046 | 0.87 | 0.39283 | 0.48 | 59 |
| CBN0048 | 0.87 | 0.78032 | 0.16 | 12 |
| CBN0520 | 0.87 | 0.36278 | 0.18 | 42 |
| CBN1404 | 0.88 | 0.61015 | 0.22 | 62 |
| CBN0559 | 0.88 | 0.8052 | 0.16 | 44 |
| CBN1379 | 0.89 | 0.39916 | 0.42 | 49 |
| CBN1594 | 0.89 | 0.26515 | 0.54 | 5 |
| CBN3446 | 0.9 | 0.71575 | 0.18 | 46 |
| CBN0582 | 0.91 | 0.05394 | 0.0 | 64 |
| CBN1602 | 0.91 | 0.55928 | 0.18 | 39 |
| CBN0443 | 0.92 | 0.3045 | 0.56 | 34 |
| CBN1096 | 0.92 | 0.07802 | 0.0 | 24 |
| CBN0013 | 0.93 | 0.01654 | 0.0 | 5 |
| CBN0051 | 0.94 | 0.40696 | 0.38 | 21 |
| CBN0038 | 0.95 | 0.33592 | 0.06 | 55 |
| CBN1258 | 0.96 | 0.91225 | 0.18 | 64 |

We observe that for each vessel, the maximum *H*_2_ persistences are close to zero in most segments. When a segment is not near the stenosis, the variation of the flow directions in this segment after preprocessing is reduced, i.e., the point cloud on *S*^2^ concentrates near one point, hence no significant two-dimensional topological structure in it. Figures 7 and 8 show the maximum *H*_1_ and *H*_2_ persistences in each segment for a few vessels as an example. The *y*-axis of all top images is the maximum *H*_1_ persistence, the *y*-axis of all images on the bottom is the maximum *H*_2_ persistence, and the *x*-axis of all images is the segment number. We can see that the vessels with ID CBN0013, CBN0582 and CBN1096 in Figure 7, which have FFR values 0.93, 0.91 and 0.92, respectively, implying a healthy state, have close-to-zero maximum *H*_2_ persistences in all segments. However, for the vessels with ID CBN0018, CBN0045 and CBN2122, which have FFR values 0.74, 0.73 and 0.66 in Figure 8, there are segments in which the maximum *H*_2_ persistences are larger than 0.3.

**Fig 7.**
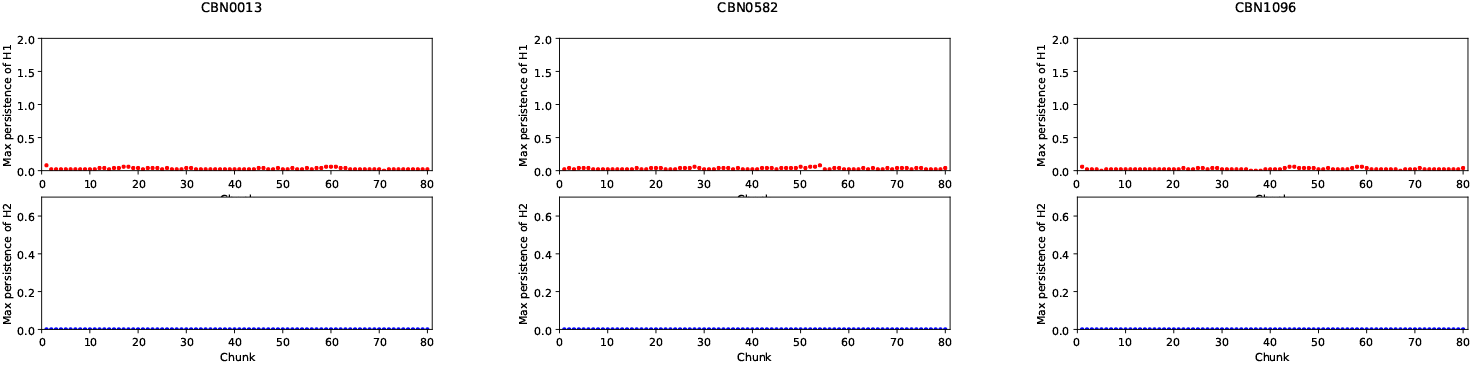
Maximum *H*_1_ (Top) and *H*_2_ (Bottom) persistences in each segment for vessels CBN0013, CBN0582, CBN1096

**Fig 8.**
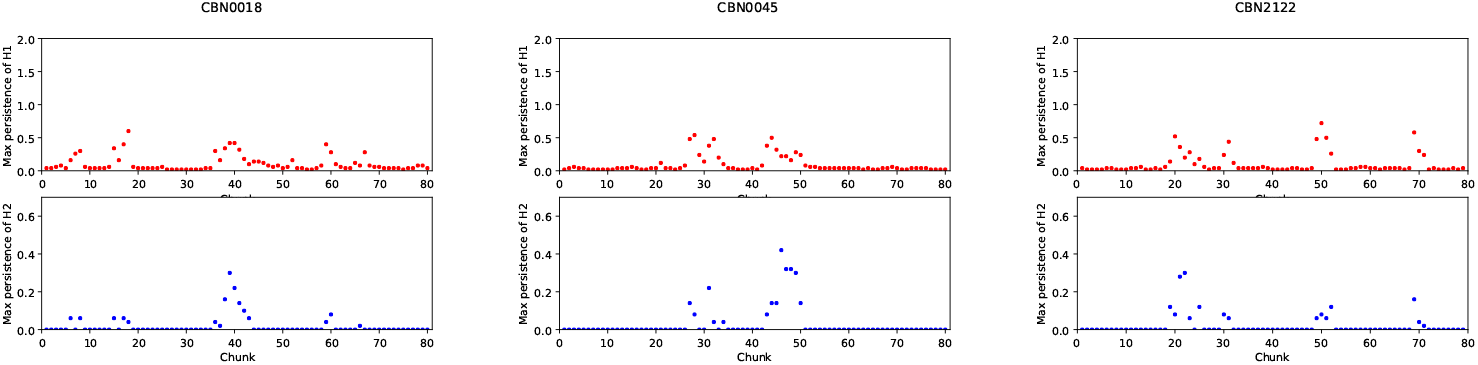
Maximum *H*_1_ (Top) and *H*_2_ (Bottom) persistences in each segment for vessels CBN0018, CBN0045, CBN2122

Figures 9 and 10 show the *H*_0_ (top figure), *H*_1_ (middle) and *H*_2_ (bottom) barcodes for the corresponding vessels in Figures 7 and 8, respectively. The barcodes are calculated using the point cloud collected from the whole vessel domains after the preprocessing. As shown in the figure, the *H*_2_ persistences for the vessels of CBN0013, CBN0582 and CBN1096 are non-existent as shown in Figure 7 while the *H*_2_ persistences for the vessels of CBN0018, CBN0045 and CBN2122 are significant as shown in Figure 8. We also observe that the *H*_1_ persistences of the vessels of CBN0013, CBN0582 and CBN1096 are smaller than those of the vessels of CBN0018, CBN0045 and CBN2122. Note that *H*_1_ persistences in Figure 9 are existent while those in Figure 7 are not. This is due the fact that the *H*_1_ in Figure 7 is only from a segment of the given vessel but the *H*_1_ in Figure 9 is calculated using the point cloud from the whole vessel.

**Fig 9.**
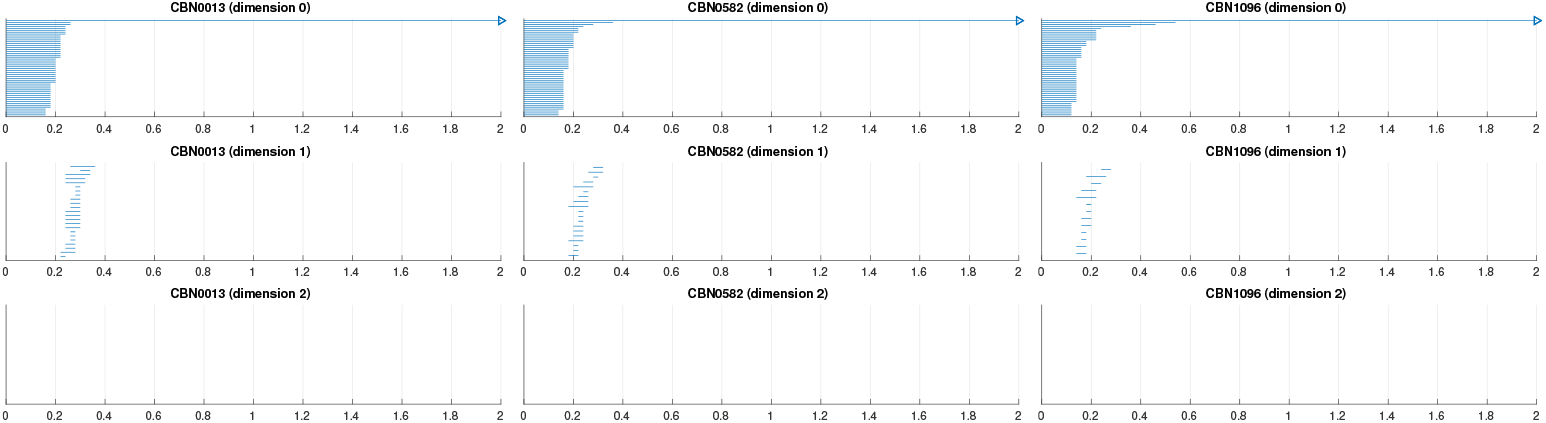
Barcodes for CBN0013, CBN0582, CBN1096 for H0 (top), H1 (middle) and H2 (bottom).

**Fig 10.**
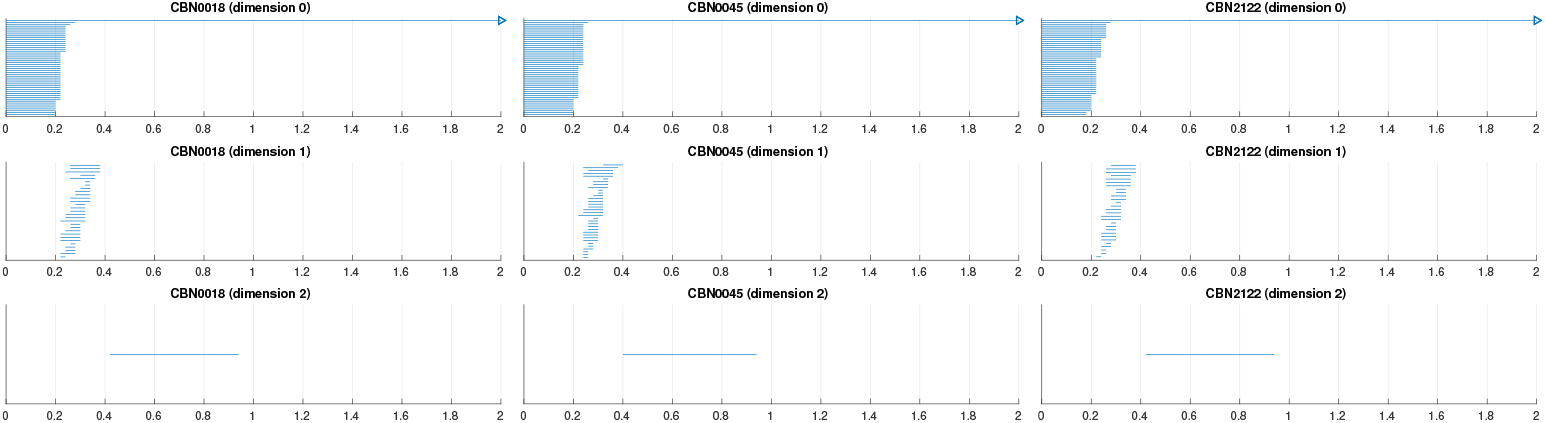
Barcodes for CBN0018, CBN0045, CBN2122 for *H*_0_ (top), *H*_1_ (middle) and *H*_2_ (bottom).

In Figure 11, we show the correlation between the FFR values and the maximum *H*_1_ persistences from Table 1. In both images, the *x*-axis is the FFR value and the y-axis is the maximum *H*_1_ persistence in the segment where the largest maximum *H*_2_ persistence occurs for each vessel. The straight lines are fitting lines after applying linear regression with least squares method to the corresponding points in each image. We can see that the *p*-values for the linear fit are 0.138 and 0.00248 for the non-preprocessed and preprocessed data, respectively. This demonstrates that we do not have a statistically significant correlation between FFR and our potential diagnostic index without the preprocessing scheme, but have a potentially useful diagnostic index with TDA if the proper preprocessing is applied.

**Fig 11.**
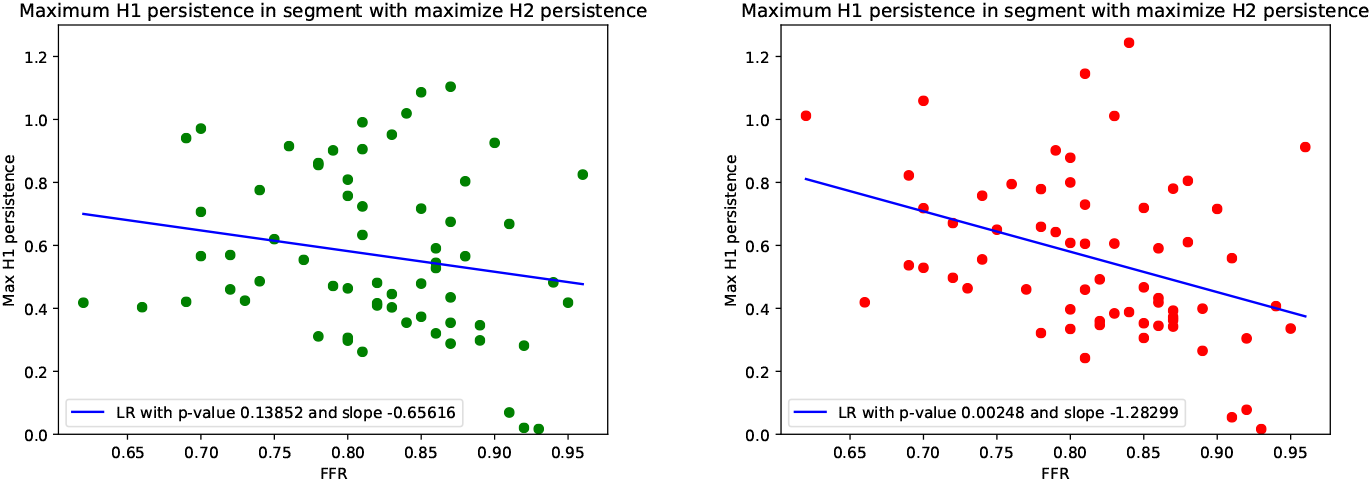
Correlation with FFR. Correlation between maximum *H*_1_ persistence in the segment where the largest maximum *H*_2_ persistence occurs and the FFR values for both before (Left) and after (Right) preprocessing with the centerline projection scheme.

**Fig 12.**
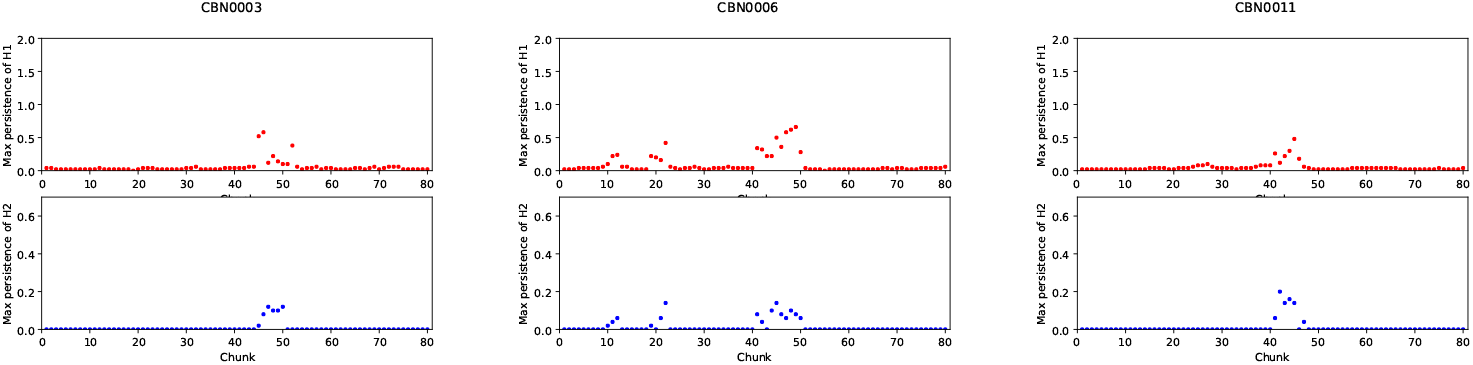
Maximum persistences for CBN0003, CBN0006, CBN0011

**Fig 13.**
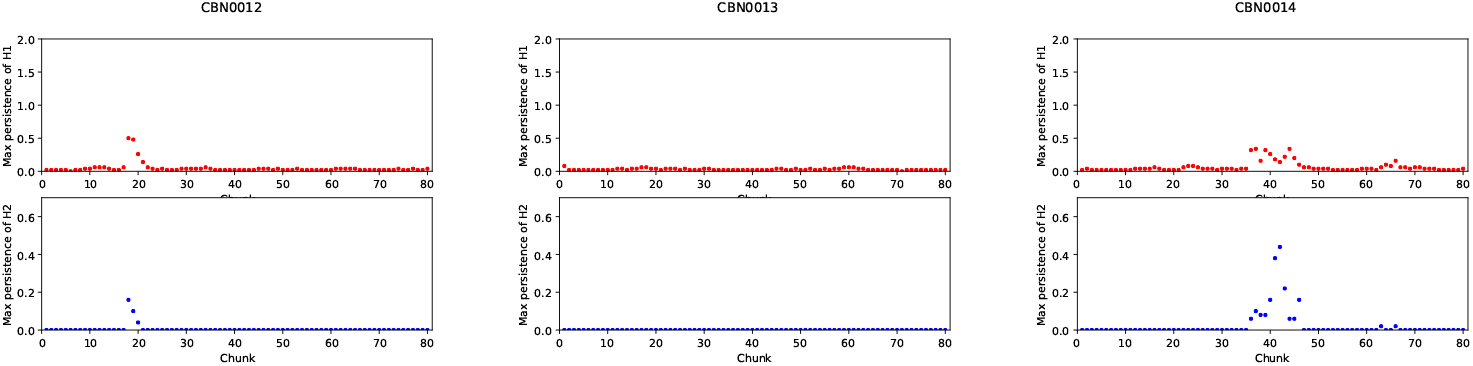
Maximum persistences for CBN0012, CBN0013, CBN0014

**Fig 14.**
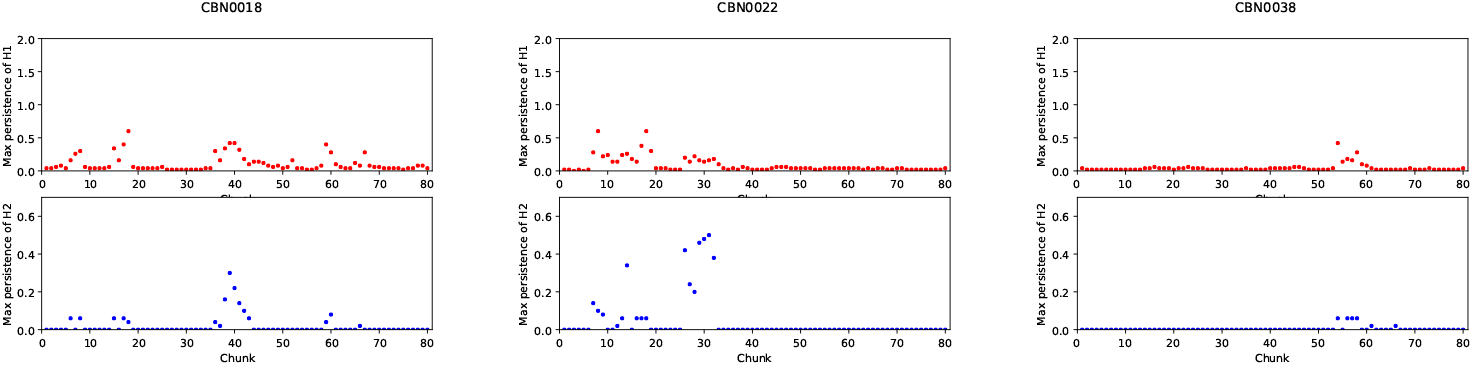
Maximum persistences for CBN0018, CBN0022, CBN0038

**Fig 15.**
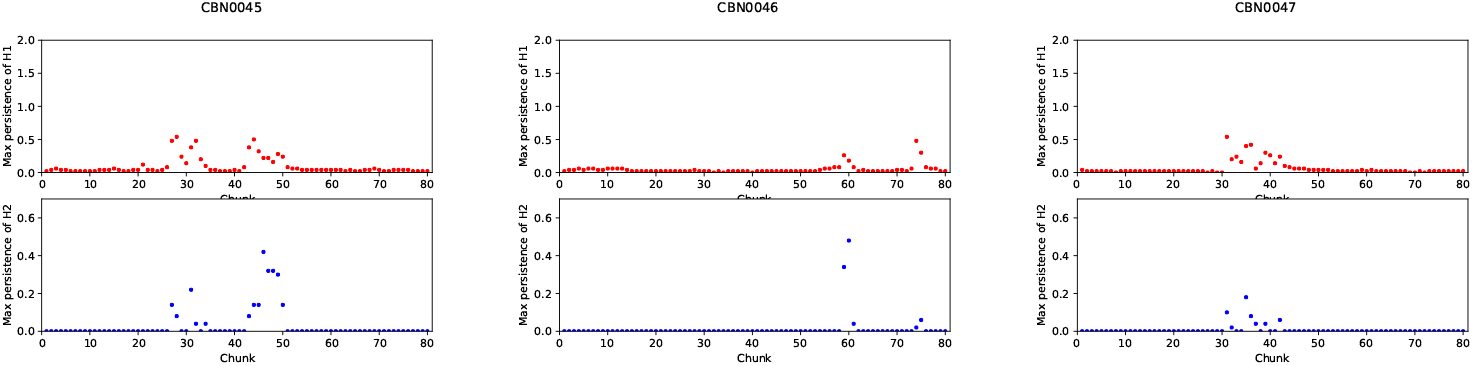
Maximum persistences for CBN0045, CBN0046, CBN0047

**Fig 16.**
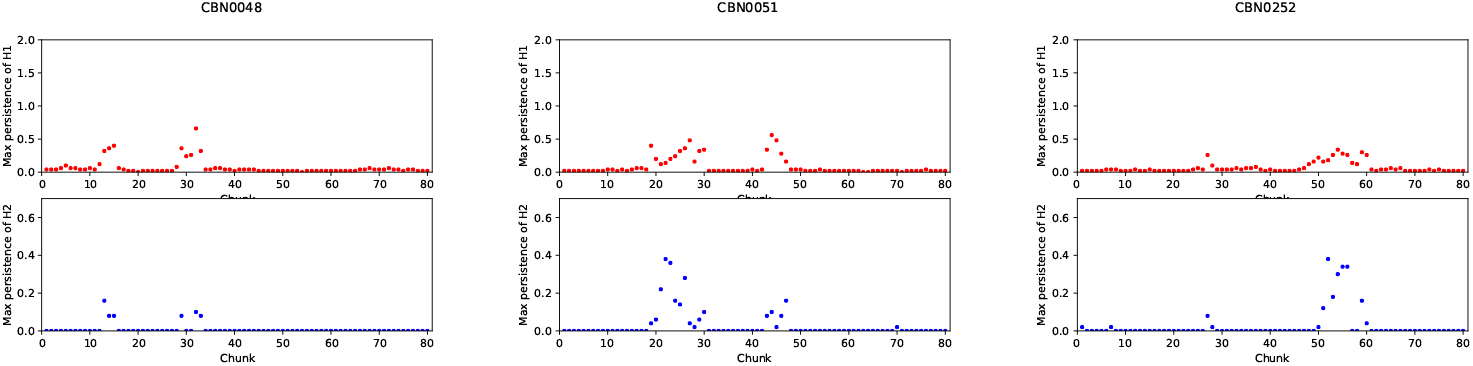
Maximum persistences for CBN0048, CBN0051, CBN0252

**Fig 17.**
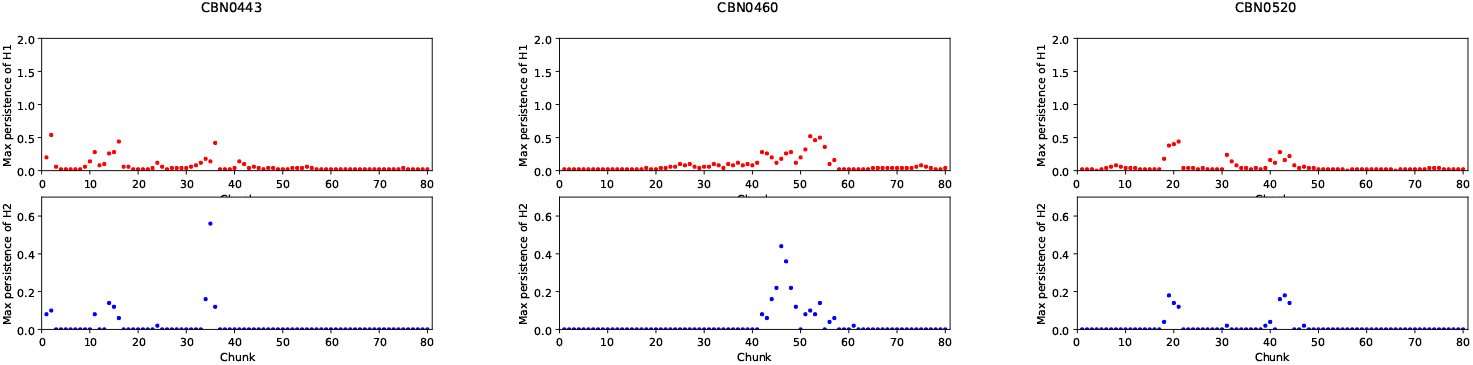
Maximum persistences for CBN0443, CBN0460, CBN0520

**Fig 18.**
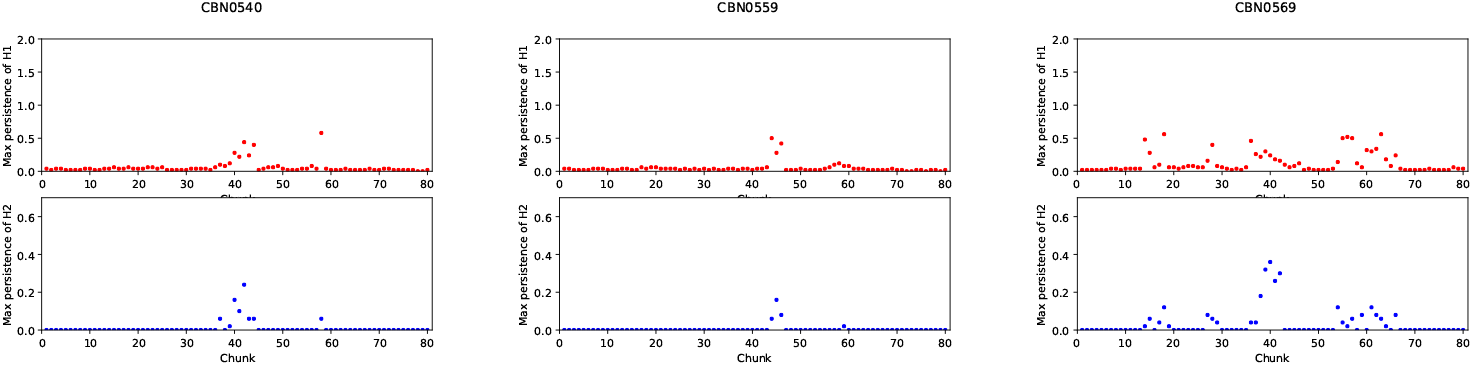
Maximum persistences for CBN0540, CBN0559, CBN0569

**Fig 19.**
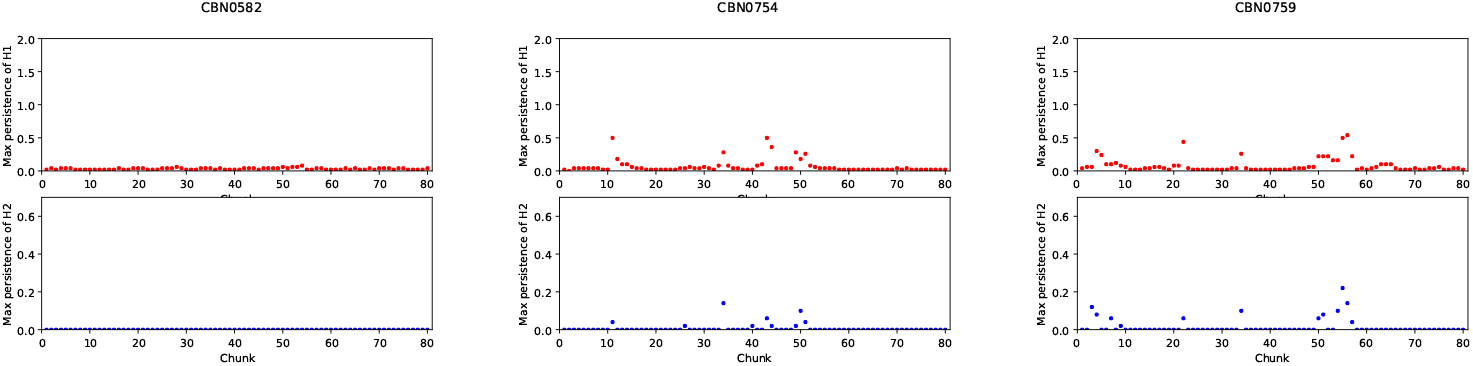
Maximum persistences for CBN0582, CBN0754, CBN0759

**Fig 20.**
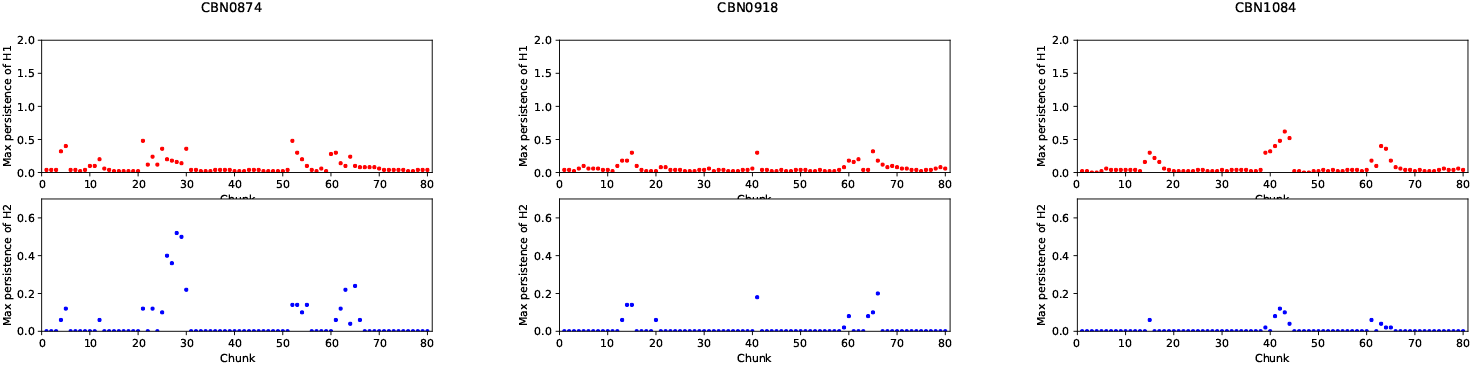
Maximum persistences for CBN0874, CBN0918, CBN1084

**Fig 21.**
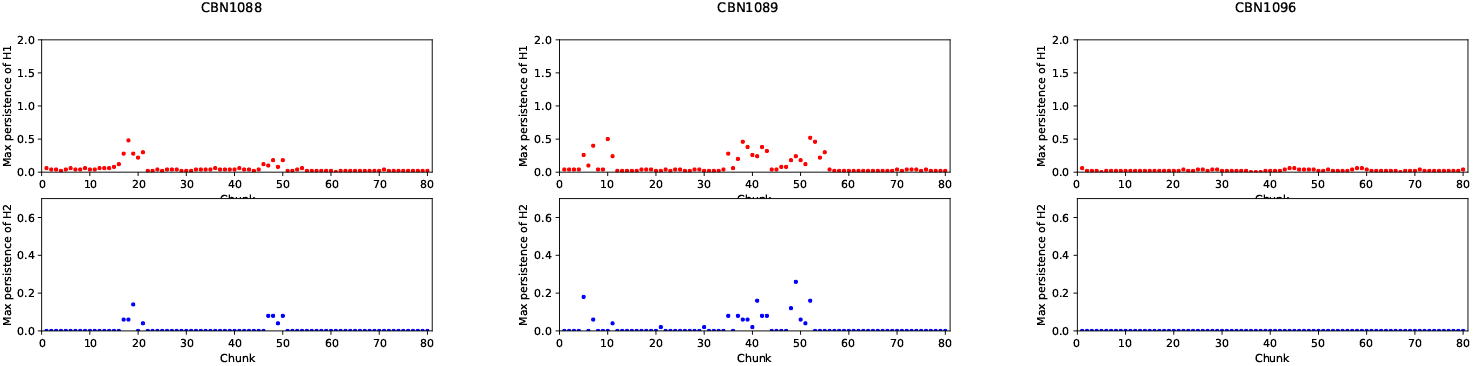
Maximum persistences for CBN1088, CBN1089, CBN1096

**Fig 22.**
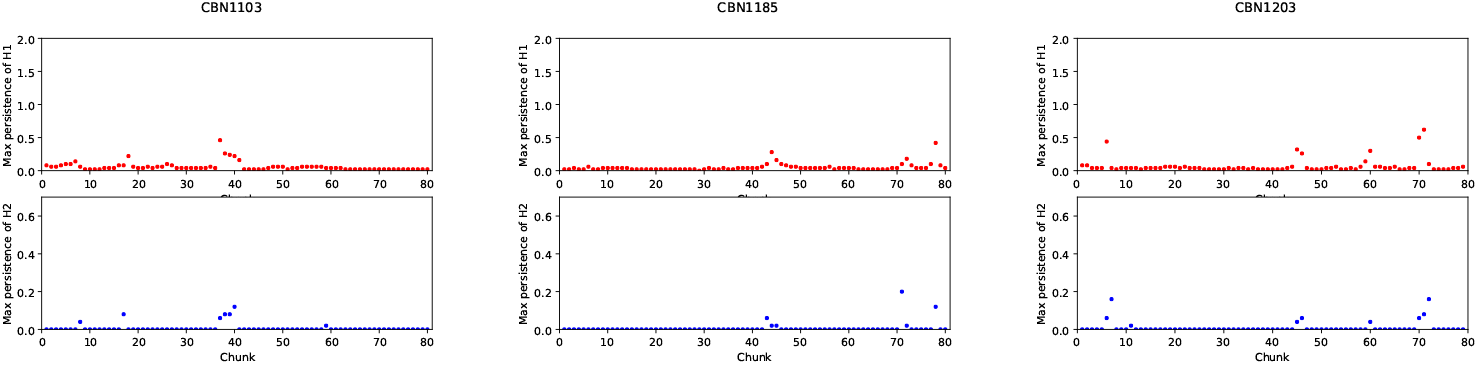
Maximum persistences for CBN1103, CBN1185, CBN1203

**Fig 23.**
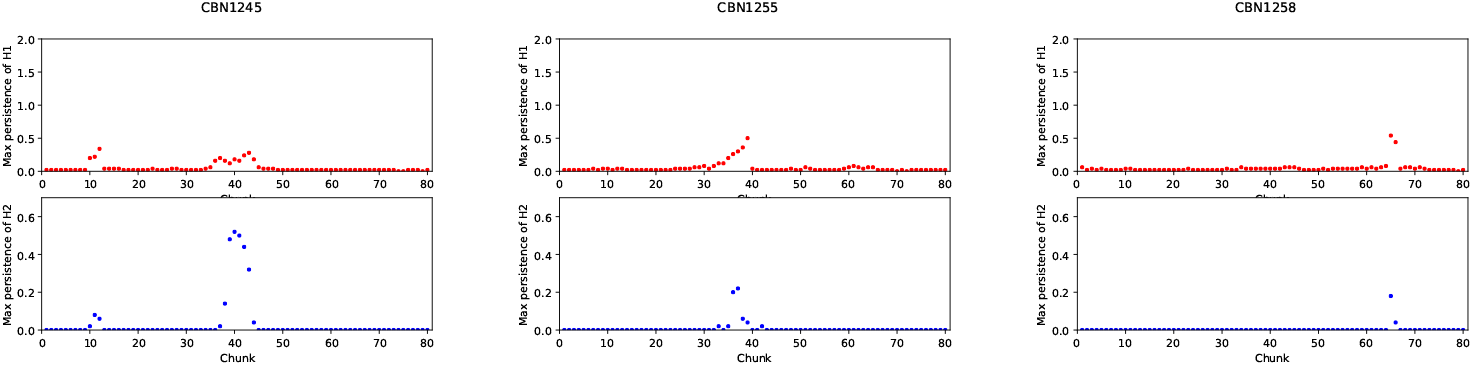
Maximum persistences for CBN1245, CBN1255, CBN1258

**Fig 24.**
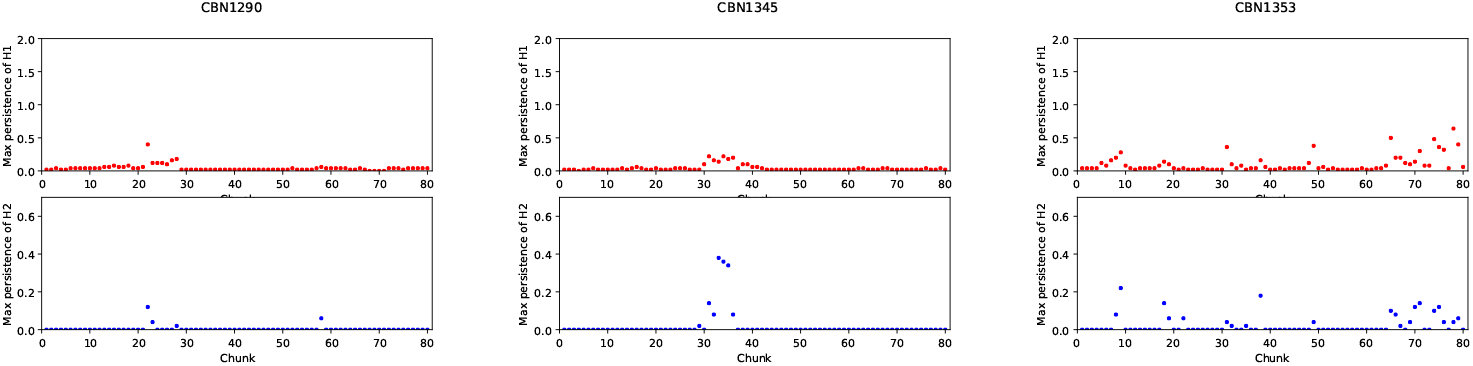
Maximum persistences for CBN1290, CBN1345, CBN1353

**Fig 25.**
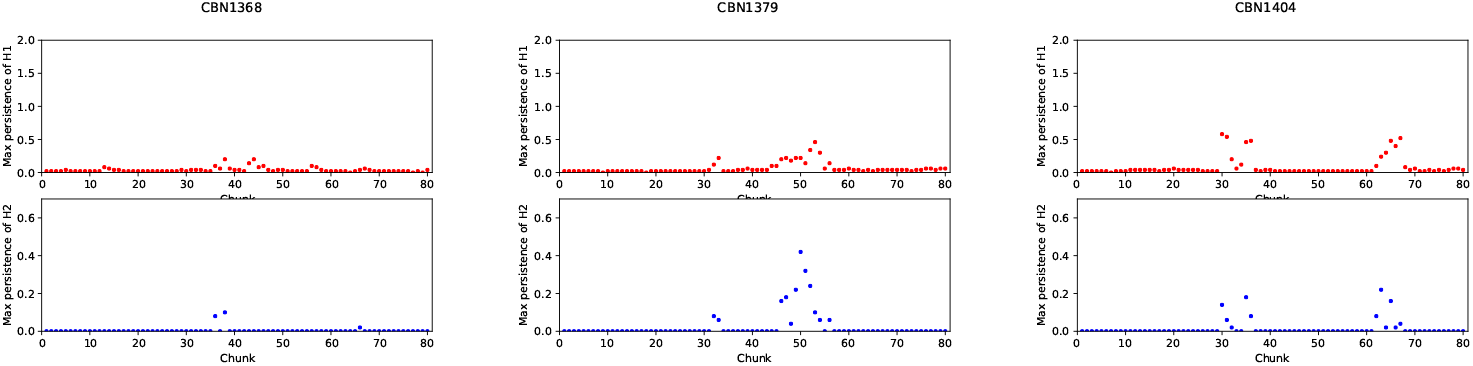
Maximum persistences for CBN1368, CBN1379, CBN1404

**Fig 26.**
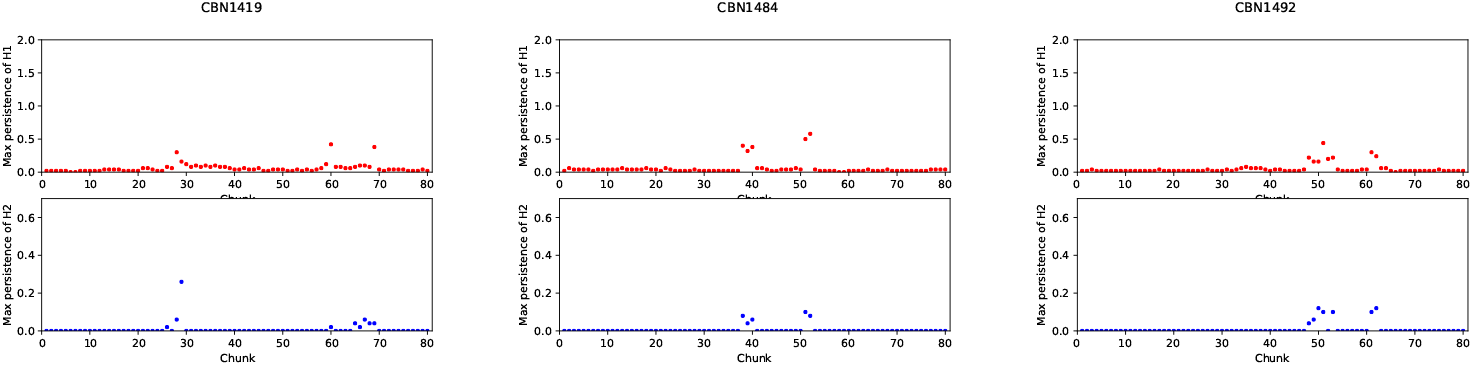
Maximum persistences for CBN1419, CBN1484, CBN1492

**Fig 27.**
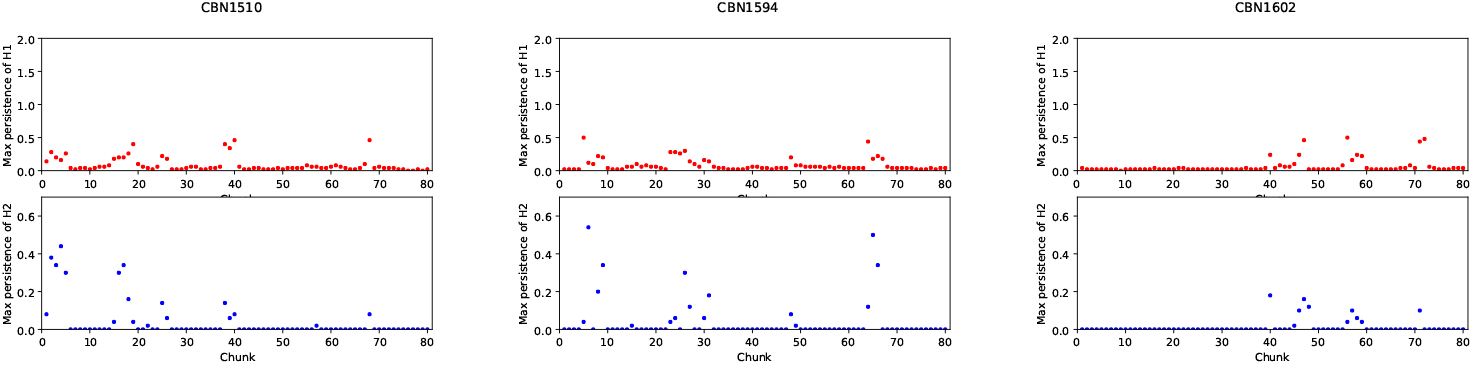
Maximum persistences for CBN1510, CBN1594, CBN1602

**Fig 28.**
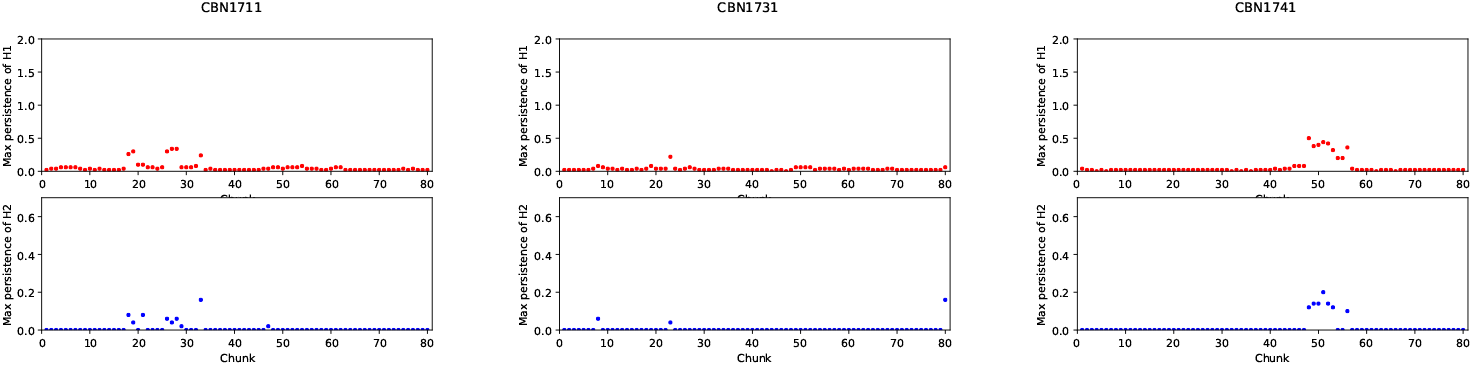
Maximum persistences for CBN1711, CBN1731, CBN1741

**Fig 29.**
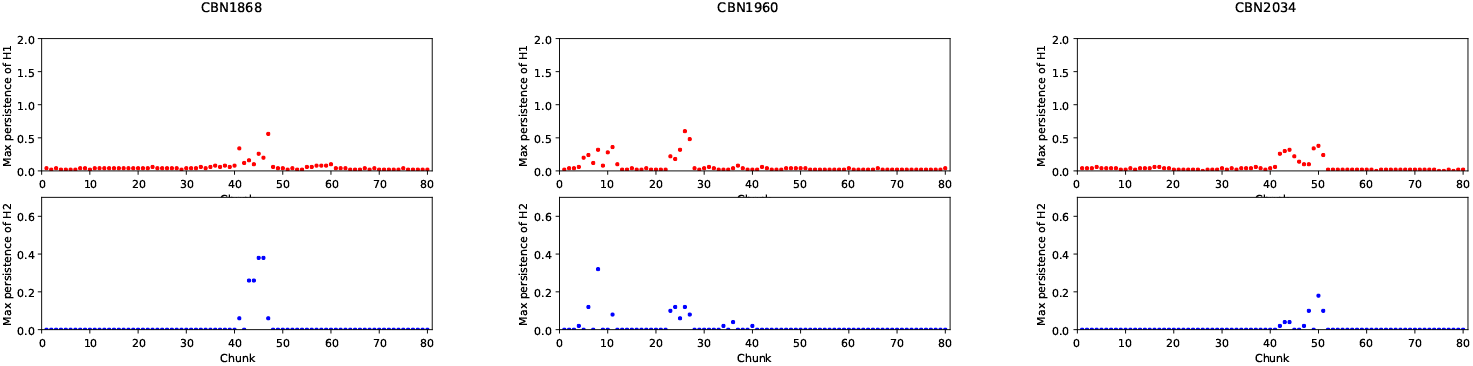
Maximum persistences for CBN1868, CBN1960, CBN2034

**Fig 30.**
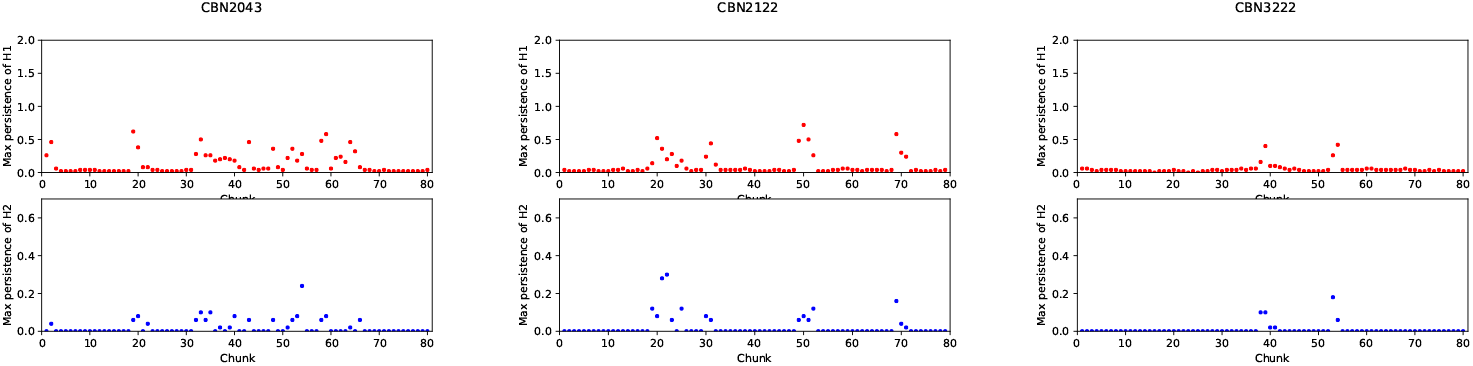
Maximum persistences for CBN2043, CBN2122, CBN3222

**Fig 31.**
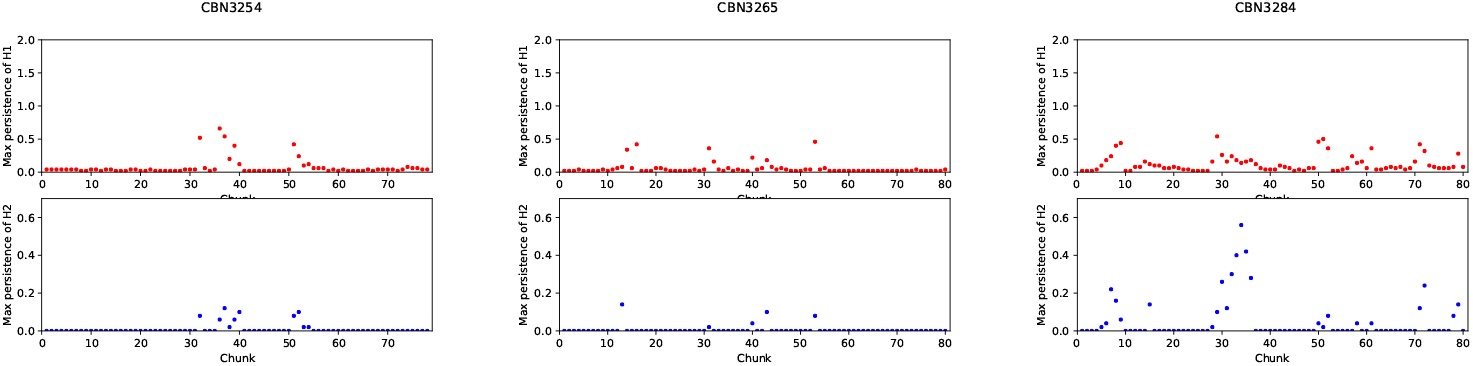
Maximum persistences for CBN3254, CBN3265, CBN3284

**Fig 32.**
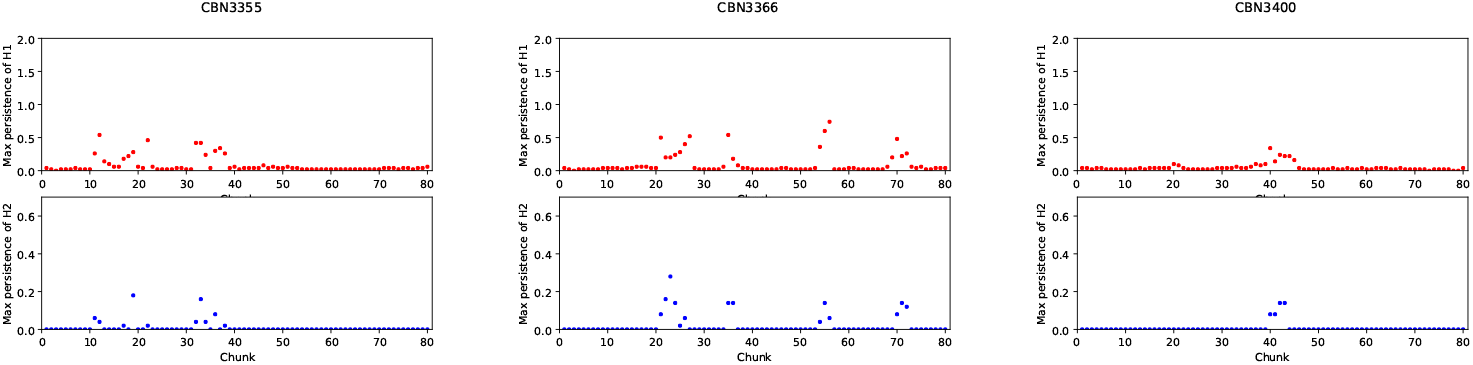
Maximum persistences for CBN3355, CBN3366, CBN3400

**Fig 33.**
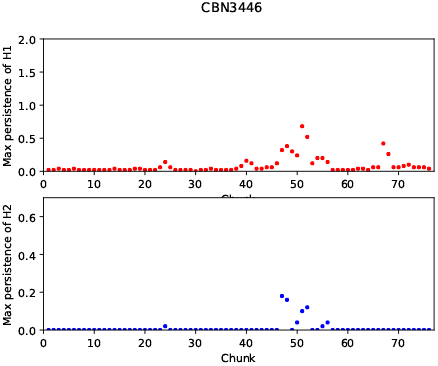
Maximum persistences for CBN3446

## Conclusion

Vascular disease is the leading cause of death worldwide. Accurate measurement of the degree of vascular disease is crucial for diagnosis and treatment. In [1] a new method based on TDA has been proposed, employing topological features as a potential diagnostic index. The proposed method was tested with the CFD solutions calculated in the straight stenotic vessels without curvatures in [1]. In this paper, we developed a preprocessing method that enables the TDA method to be applied to the curved stenotic vessel. We applied the proposed method to clinically sourced data, showing a statistically significant correlation to the gold standard diagnostic index, the FFR. It was also shown that the preprocessing method was necessary to achieve a statistically significant correlation. This validates the proposed method as a potentially powerful diagnostic tool.

In future work we plan to refine the preprocessing method and make adaptations for application to more general vascular geometries such as bifurcated vessels, multiple stenoses. There is also potential for improvement of computational efficiency, parameter selection and other methodologies in the preprocessing scheme.

## S1 Appendix

**S1 Figures: Maximum** *H*_1_ **and***H*_2_ **versus segment number** In this appendix, we provide the maximum *H*_1_ and *H*_2_ persistences calculated in each vessel segment used in preprocessing steps versus the segment number for those vessels in Table 1. These figures show not only the flow complexity but also the possible locations of stenosis.

## Acknowledgments

This work has been supported by Samsung Science & Technology Foundation under grant number SSTF-BA1802-02. Christopher Bresten is supported by KRF program by National Research Foundation under NRF-2018H1D3A1A01074679.

